# The role of dopaminergic nuclei in predicting and experiencing gains and losses: A 7T human fMRI study

**DOI:** 10.1101/732560

**Authors:** Laura Fontanesi, Sebastian Gluth, Jörg Rieskamp, Birte U. Forstmann

**Affiliations:** University of Basel; University of Amsterdam

**Keywords:** reward, punishment, midbrain, substantia nigra, ventral tegmental area

## Abstract

The ability to predict the outcomes of actions based on experience is crucial for making successful decisions in new or dynamic environments. In animal studies using electrophysiology, it was found that dopamine neurons, located in the substantia nigra (SN) and the ventral tegmental area (VTA), have a crucial role in feedback-based learning. However, human neuroimaging studies have provided inconclusive results. The present work used ultrahigh field (7 Tesla) structural and functional MRI and optimized protocols to extract SN and VTA signals in human participants. In a number-guessing task, we found significant correlations with reward prediction error and risk in both the SN and the VTA and no correlation with expected value. We also found a surprise signal in the SN. These results are in line with a recent framework that proposed a differential role for the VTA and the SN in, respectively, learning of values and surprise.

## Introduction

In order to adapt to an ever-changing environment, it is crucial for individuals to correctly predict the outcomes of their choices, as well as to update their expectations when they happen to be wrong. These learning processes were formalized within the reinforcement learning (RL) framework (Sutton & Barto, 1998), unifying the fields of psychology and artificial intelligence. In this framework, the reward prediction error (RPE) is defined as the difference between the expectations and the experienced rewards or punishments, and guides learning: New expectations are a weighted sum of past expectations and the RPE. By presenting participants in the lab with different options and providing feedback after every decision, psychologists and neuroscientists can investigate the cognitive processes related to *expectations* and *feedback processing*. Expectations can be separated into the expected value (EV), which can be defined as the mean expected outcome, and risk, which is often defined as the expected variance of the outcomes (Markowitz, 1952). Feedback-related processes are the deviation from previous expectations (i.e., the RPE) and the salience of the outcome (i.e., surprise, see Methods section).

A highly distributed network related to expectations and feedback processing was found in both the animal and the human brain. Electrophysiological studies in rodents and non-human primates showed that midbrain dopaminergic neurons (i.e., in the substantia nigra, SN – specifically in its pars compacta, SNc – and in the ventral tegmental area, VTA) fire more, equal, or less in association with a positive, zero, or negative RPE (Bayer & Glimcher, 2005; Schultz, 1998, 2015), respectively, and their firing ramps up faster with increasing risk expectations (Fiorillo, Tobler, & Schultz, 2003). Firing of cells in the SNc has also been associated with surprise (Matsumoto & Hikosaka, 2009). Because dopamine nuclei are more challenging to target using non-invasive neuroimaging techniques, studies using human participants mainly focused on dopamine target areas (Arias-Carrión, Stamelou, Murillo-Rodríguez, Menéndez-Gonzáles, & Pöppel, 2010). Neural correlates of the RPE have been found in the ventral striatum and an expected reward signal has been found in ventral striatum, amygdala, as well as in frontal areas such as the orbital frontal cortex and the ventromedial prefrontal cortex (for an overview see, e.g., Bartra, McGuire, & Kable, 2013; Clithero & Rangel, 2014; O’Doherty & Bossaerts, 2008). Both ventral striatum and anterior insula were found to signal predicted risk and surprise (Fouragnan, Retzler, & Philiastides, 2018; Preuschoff, Bossaerts, & Quartz, 2006; Singer, Critchley, & Preuschoff, 2009).

The measurement of small dopaminergic nuclei signaling using fMRI is very challenging. One challenge pertains to the higher concentration of iron in the SN (Drayer et al., 1986). This high concentration causes differences in the magnetic properties of the SN compared to, for example, cortical areas, and asks for customized structural and functional MRI scanning protocols (e.g., reduced echo times). Another problem is physiological noise affecting the fMRI data due to the proximity of these areas to major arteries and cerebrospinal fluid. Finally, their limited volume and distance from the receiving elements of the scanner, combined with anatomical variability and standard procedures such as spatial smoothing, lead to a high risk of mixing signals from neighboring nuclei (de Hollander, Keuken, & Forstmann, 2015; de Hollander, Keuken, van der Zwaag, Forstmann, & Trampel, 2017; Eapen, Zald, Gatenby, Ding, & Gore, 2011; Forstmann, de Hollander, van Maanen, Alkemade, & Keuken, 2017).

Because of these challenges, only very few neuroimaging studies have directly measured activation of small dopaminergic nuclei in human participants. Furthermore, these studies reported contradicting evidence. D’Ardenne, McClure, Nystrom, and Cohen (2008) found positive but not negative RPE in the VTA. Pauli et al. (2015) found only a positive RPE in the SNc, a negative RPE in the pars reticulata of the SN (SNr), as well as a negative expected value signal in the SNr. Zhang, Larcher, Misic, and Dagher (2017) found that, while the medial part of the SN encoded RPE, the lateral and ventral parts encoded surprise.

To the best of our knowledge, previous studies with human subjects (1) have not compared the signal of the VTA and the SN (except D’Ardenne et al. (2008)), (2) have not looked at the variables related to expectations and feedback processing altogether (i.e., they did not always include EV, risk, RPE, and surprise); (3) have not addressed the above-mentioned fMRI-specific challenges. In particular, previous studies have used high-field 3 Tesla (3T) MRI, spatial smoothing, and did not draw individual masks to delineate the VTA or the SN, but relied instead on group-based coordinates or atlases.

Ultra-high-field (UHF) 7 Tesla (7T) MRI can help to increase signal-to-noise ratio (SNR) and BOLD contrast-to-noise ratio (CNR), leading to a more refined spatial resolution without loss of power or need for spatial smoothing (van der Zwaag, Schäfer, Marques, Turner, & Trampel, 2015). In the present study, we used UHF-fMRI in combination with scanning protocols tailored to extract signals from subject-specific masks of the midbrain to overcome some of the previous limitations and clarify the findings of previous studies, especially regarding the function of the VTA and the SN (Trutti, Mulder, Hommel, & Forstmann, 2019). By adapting the number-guessing paradigm proposed by Preuschoff et al. (2006), we also investigated important variables such as risk and surprise, as well as EV and RPE, thereby targeting processes of both expectation and feedback processing.

## Results

To investigate the role of the VTA and the SN in expectation and feedback processing, we tested participants in a number-guessing task (Figure 1) in a MRI session. In this task, there are three main events per trial. First, participants have to predict whether the first or the second of two numbers (between 1 and 5) will be higher: this prediction corresponds to their initial bet, as if the prediction is correct they will win 5 euros and if the prediction is incorrect they will lose 5 euros. Then, they are shown the first of the two numbers, which changes the EV and risk of the choice options. Finally, participants are shown the second number, together with the reward, which is associated with a specific RPE and surprise, depending on the initial bet and on the first number. Participants were also invited to a separate MRI session, in which multimodal, high-resolution anatomical images were acquired (Figure 2). This procedure allowed us to identify the region of interests (ROIs) at an individual level and to then extract the signal from each ROI to test for correlations with EV, risk, RPE, and surprise.

**Figure 1.**
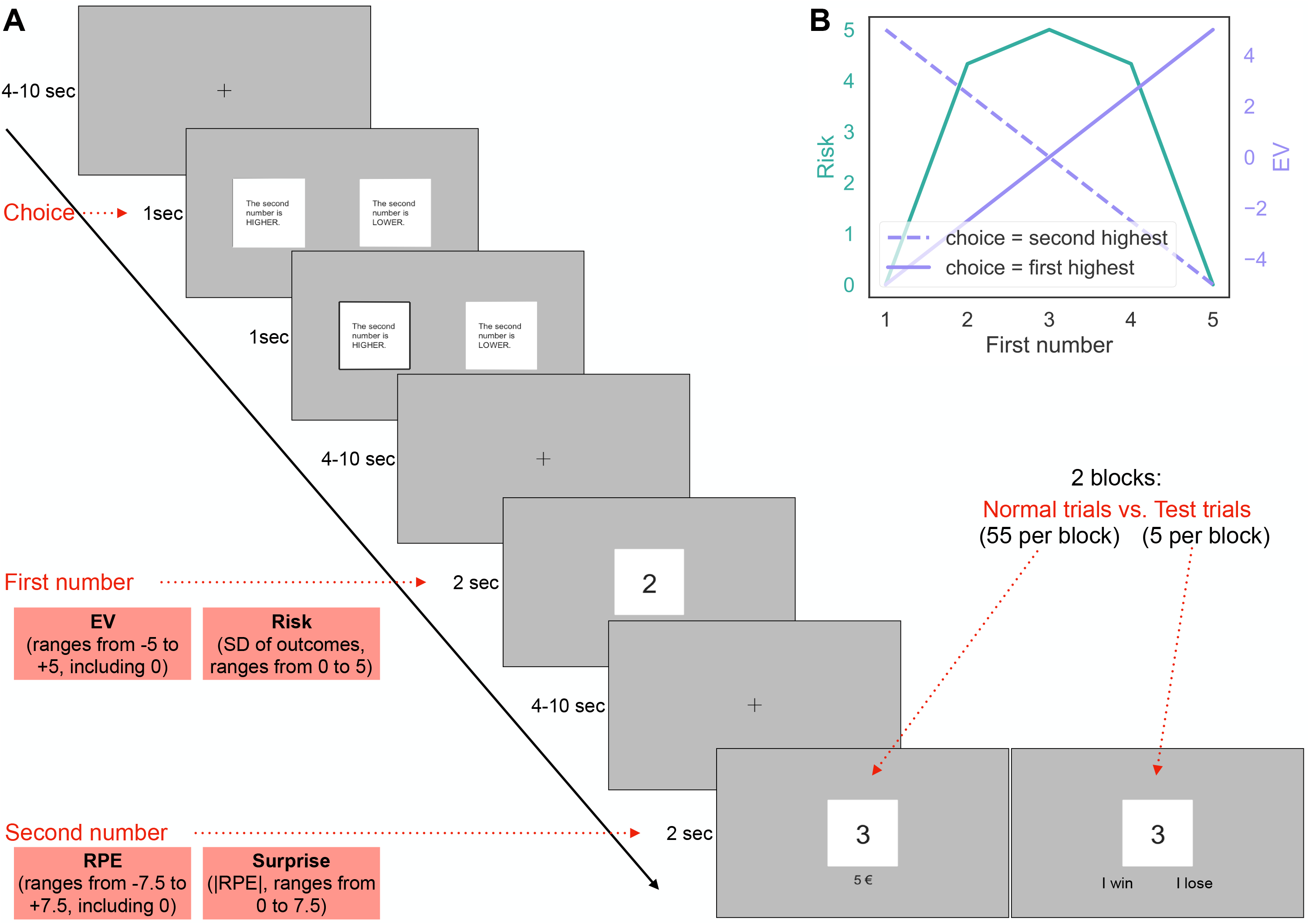
Experimental design. A. Example of a single trial. Between each event and at the beginning of each trial, a fixation cross is presented for a period of time between 4 and 10 seconds. A bet has to be placed within 1 second, and a rectangle is drawn around the corresponding choice for 1 more second. The first number is then shown for 2 seconds: In this example, the expected reward is 2.5 euros, and the risk is 4.3. Finally, the second number is shown for 2 seconds: In this case, both the reward prediction error and the surprise are 2.5. In test trials (approximately 8%) participants have to specify whether they won or lost. B. Relationship between risk and expected reward when the first number is shown, depending on the choice.

**Figure 2.**
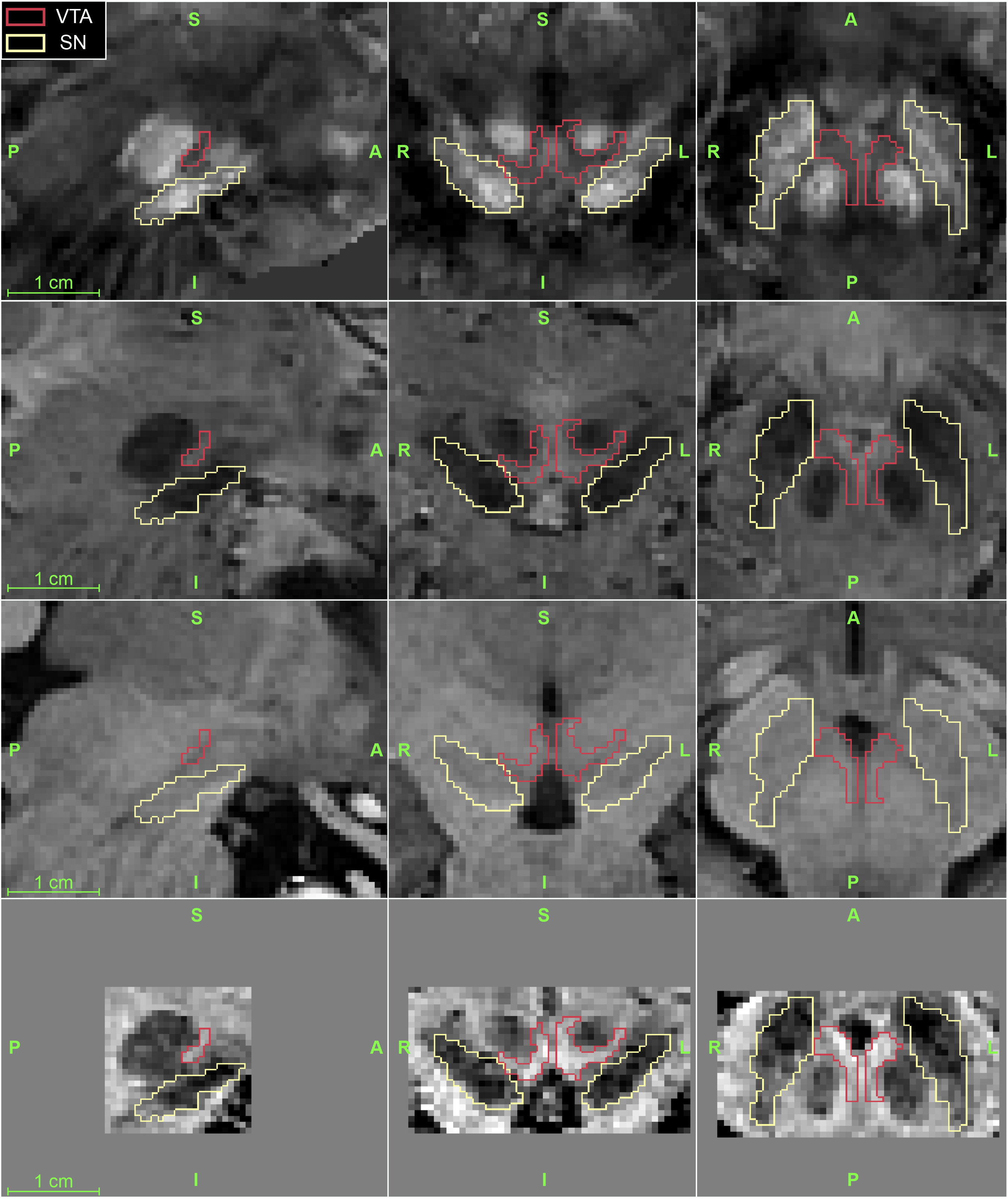
Detail of the midbrain area of one participant in the sagittal (first column), coronal (second column), and axial (third column) planes. The first row is the QSM image, used for SN segmentation. The second and third row are, respectively, the average between the third and fourth echo of the 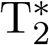-weighted, and the T_1_-weighted images. To obtain the image in fourth row, the images in the second and third row were normalized within the midbrain area (the non-homogeneous grey area in the last row) and then summed. This image was used for VTA segmentation, as it shows a contrast of both iron-rich nuclei and of the CSF.

In the following sections, we report the behavioral results of the card-guessing task, the results of the anatomical segmentation of the ROIs, the fMRI analyses results limited to the ROIs, as well as across the whole brain.

### Behavior

To check whether participants were engaged in the task, we introduced test trials in which, instead of revealing their reward, participants had to say whether they won or lost in that specific trial. This was only possible if they still remembered the first number and their initial bet. Three blocks (from three different participants) were discarded based on behavior: One block was discarded because three out of the five test trials were incorrect, and the other two blocks were discarded because twelve out of sixty missed bets.

In the remaining blocks, and over the two blocks (i.e., 120 total trials), participants made on average 1.0 mistakes (SD=1.05, min=0, max=4), missed on average 4.48 trials (SD=3.65, min=0, max=12), and chose on average the right option on 57.81 trials (SD=13.75, min=21, max=88).

### Anatomical masks

To measure the inter-rater reliability of the individual SN and VTA segmentation, we calculated Dice Scores (see Table 1). In general, higher scores were obtained for the SN as compared to the VTA. This is not surprising, because Dice scores are sensitive to overall size (the SN is approximately 3.7 times bigger than the VTA), and because the VTA lacks clear anatomical borders. By only keeping those voxels that both raters agreed on (i.e., the conjunction masks), we ensured that the voxels included in the analyses lie exclusively in the investigated ROIs.

**Table 1.**
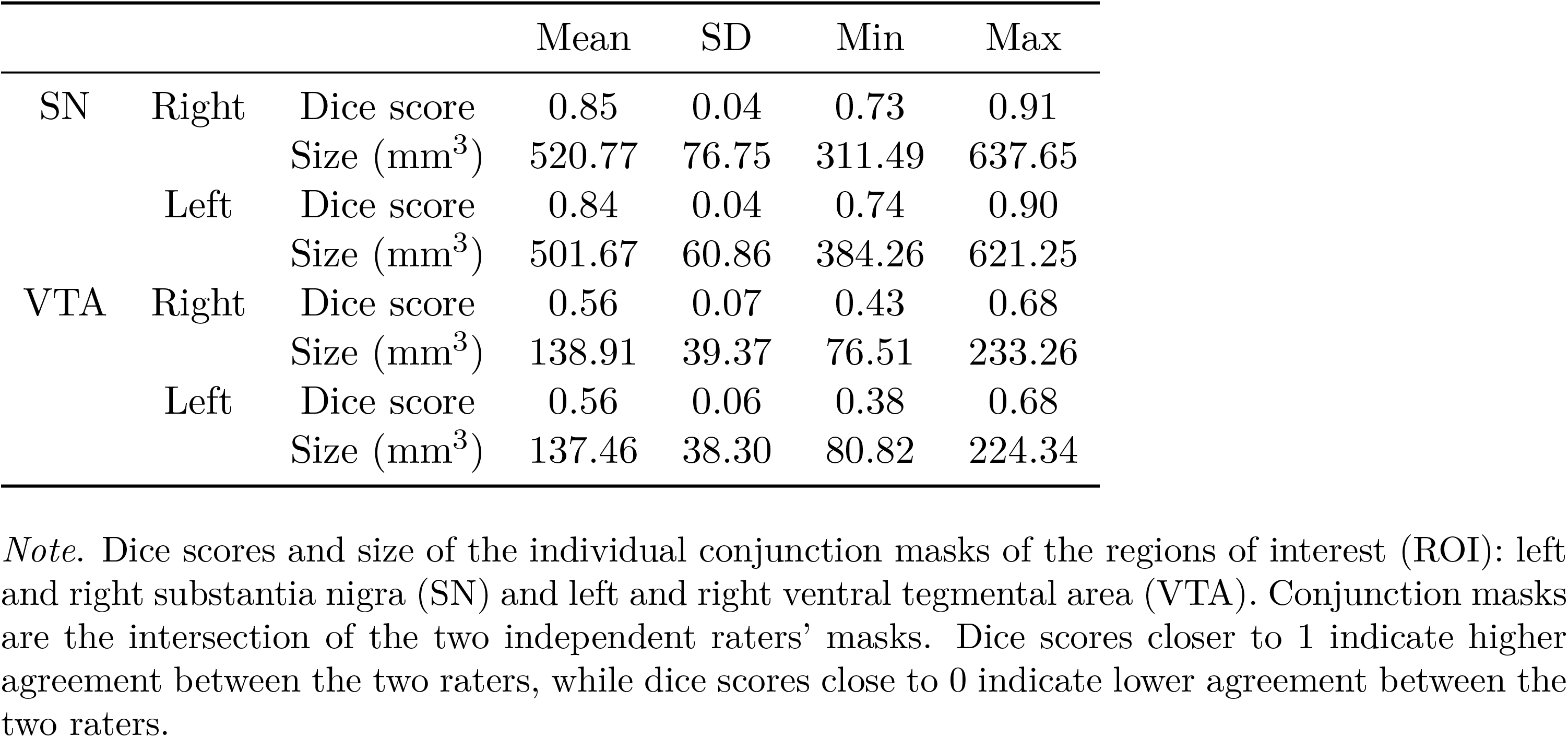
Anatomical segmentation results.

|  |  |  | Mean | SD | Min | Max |
| --- | --- | --- | --- | --- | --- | --- |
| SN | Right | Dice score | 0.85 | 0.04 | 0.73 | 0.91 |
|  |  | Size (mm<sup>3</sup>) | 520.77 | 76.75 | 311.49 | 637.65 |
|  | Left | Dice score | 0.84 | 0.04 | 0.74 | 0.90 |
|  |  | Size (mm<sup>3</sup>) | 501.67 | 60.86 | 384.26 | 621.25 |
| VTA | Right | Dice score | 0.56 | 0.07 | 0.43 | 0.68 |
|  |  | Size (mm<sup>3</sup>) | 138.91 | 39.37 | 76.51 | 233.26 |
|  | Left | Dice score | 0.56 | 0.06 | 0.38 | 0.68 |
|  |  | Size (mm<sup>3</sup>) | 137.46 | 38.30 | 80.82 | 224.34 |

*Note.* Dice scores and size of the individual conjunction masks of the regions of interest (ROI): left and right substantia nigra (SN) and left and right ventral tegmental area (VTA). Conjunction masks are the intersection of the two independent raters' masks. Dice scores closer to 1 indicate higher agreement between the two raters, while dice scores close to 0 indicate lower agreement between the two raters.

In addition to the Dice scores, we also calculated the percentage of overlap between our individual conjunction masks and previously proposed group-level subdivisions of the SN and the VTA ^1^ (Pauli, Nili, & Tyszka, 2018; Zhang et al., 2017), transformed to the individual space (see Figure 3). This measure gives an idea of how much signal from the neighbouring nuclei is mixed with the signal of the targeted structure when using population-based instead of individual masks. This measures does not include further mixing of the signal due techniques such as spatial smoothing (which may further increase this measure). We found significant overlap between the medial parts of the SN of the group-level subdivisions and our individual VTA masks. Specifically, there was a mean overlap of 7.23 percent (SD=10.14, min=0.00, max=34.58, t(53)=5.19, p*<*0.001) with the medial part of the SNc (mSNc), and a mean overlap of 1.3 percent (SD=2.14, min=0.00, max=8.36, t(53)=4.41, p*<*0.001) with the lateral part of the SNc (lSNc) as defined by Zhang et al. (2017); and a mean overlap of 1.56 percent (SD=2.21, min=0.00, max=11.93, t(53)=5.13, p*<*0.001) with the SNc as defined by Pauli et al. (2018). We also found a significant overlap between Pauli et al. (2018)’s subdivisions of the VTA (i.e., labelled VTA and the parabrachial pigmented area or PBP, where VTA denotes the more medial and PBP denotes the more lateral part) and our individual SN masks. Specifically, there was a mean overlap of 7.76 percent (SD=9.81, min=0.00, max=58.76, t(53)=5.76, p*<*0.001) with Pauli et al. (2018)’s VTA and a mean overlap of 8.81 percent (SD=6.47, min=0.00, max=25.76, t(53)=9.91, p*<*0.001) with Pauli et al. (2018)’s PBP.

**Figure 3.**
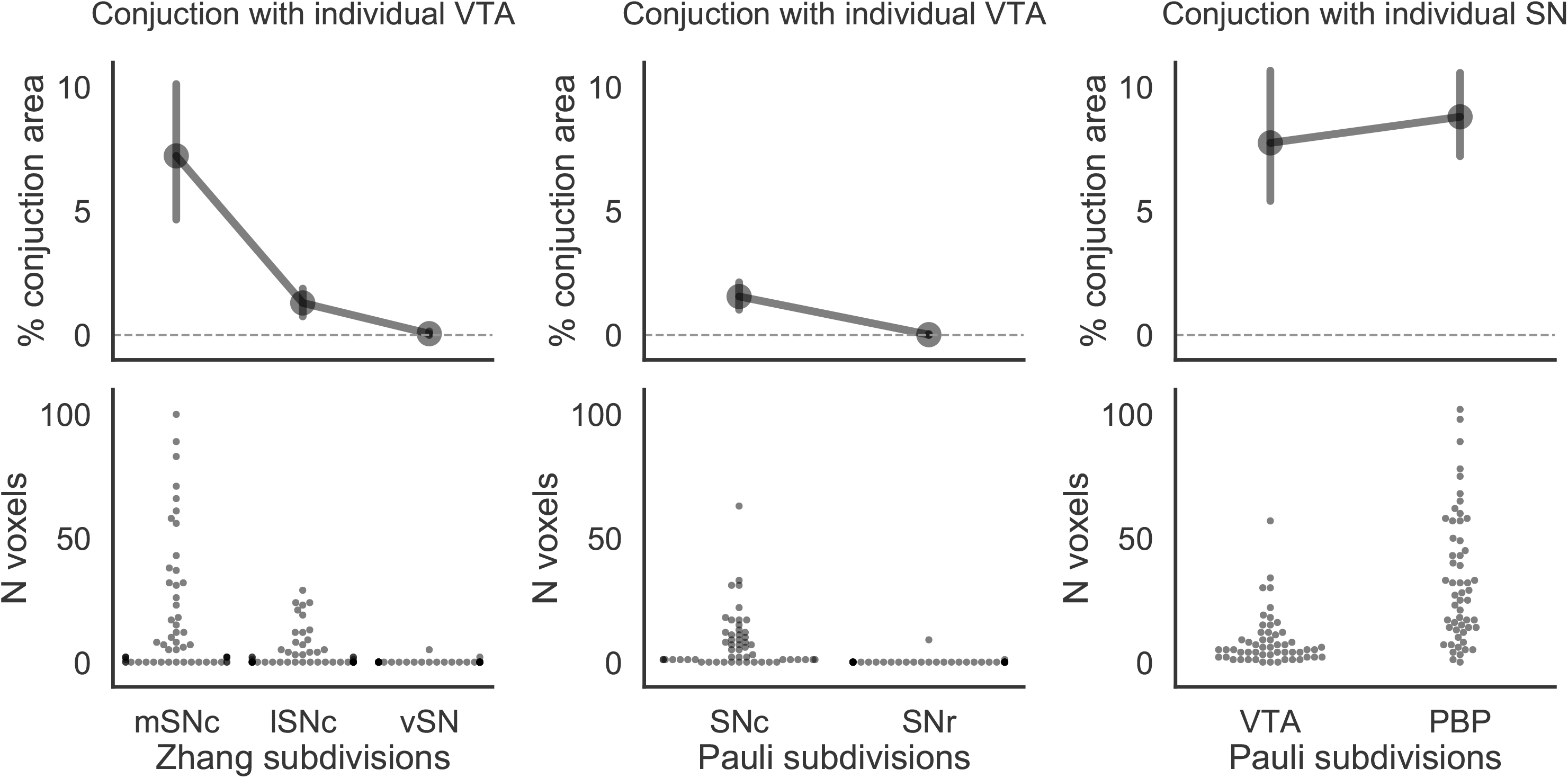
Conjunction between the population-defined substantia nigra (SN) and ventral tegmental area (VTA) subdivisions and, respectively, individually defined VTA and SN segmentation. The SN subdivisions were taken from either Zhang et al. (2017) or Pauli et al. (2018) studies, while the VTA subdivisions were taken from Pauli et al. (2018) study. Top row: percentage of the conjuction area over the subdivision area. Dots represent mean across subjects, while error bars represent 95% confidence intervals. Bottom row: swarmplot showing the number of voxels in the conjuction per subject. The medial parts of the SN (mSNc, and SNc) overlap more with the VTA than the lateral and ventral parts of the SN (lSNc, vSN, and SNr). Both the ventral (VTA) and lateral (PBP) parts of the VTA overlap with the SN.

To gain better insight into the anatomical specificity of the SN and VTA, we plotted Pauli et al. (2018)’s and Zhang et al. (2017)’s subdivisions of the SN and the VTA on the individual data using different contrasts: Figure S1, S2, and S3 show, respectively, a comparison between Pauli et al. (2018)’s atlas with our probabilistic VTA and SN maps in the MNI space, a comparison between Zhang et al. (2017)’s atlas with our probabilistic VTA and SN maps in the MNI space, and a comparison between Pauli et al. (2018)’s and Pauli et al. (2018)’s atlases in the individual space of one example subject. Although the group-level masks appear to be accurate to some extent, they often include neighbouring areas (such as the red nucleus, see the top left quadrant in Figure S3) or exclude parts of the targeted areas (such as in the lower right quadrant in Figure S3). Therefore, only by drawing individual masks and avoiding spatial smoothing, we can be sure to not mix signals from different midbrain nuclei.

Finally, we calculated the temporal signal-to-noise (tSNR) across the ROIs (see Figure S4). The tSNR was lower, yet comparable to the one reported by de Hollander et al. (2017).

### ROI-wise GLM

For the fuctional analyses, two blocks of trials (from two different participants) were discarded based on excessive head movements, having a mean framewise displacement (FD, Power et al., 2014) over .3 mm. Because one of these blocks was already discarded based on behavior, a total of four blocks was excluded from the final analyses. In the remaining blocks, and over the two blocks, participants had an average mean FD of .14 mm (SD=.06, min=.04, max=.27).

Results of the ROI-wise GLM are shown in Table 2 and Figure 4. First, we investigated the signal related to expectations (i.e., EV and risk) in both the SN and the VTA, corresponding to the presentation of the first number. We found no parametric correlations between signal in any of the ROI with the EV, with the Bayes Factor (BF) pointing to substantial (Jeffreys, 1961) evidence for the null hypothesis. However, there were significant correlations with risk in both the left-VTA (t(26)=−2.34, p*<*0.05) and the left-SN (t(26)=−2.44, p*<*0.05). Next, we investigated the signal related to feedback processing (i.e., RPE and surprise), corresponding to the presentation of the second number. There were significant correlations with RPE in the left- and right-VTA (t(26)=3.12, p*<*0.05, and t(26)=2.76, p*<*0.05) and in the right-SN (t(26)=2.54, p*<*0.05). Finally, we found a correlation with surprise in the right-SN (t(26)=2.32, p*<*0.05), and no effect in the VTA, with the BF providing substantial support for the null hypothesis. In sum, both the VTA and the SN were linked to risk before the outcome was revealed as well as to RPE after the outcome was revealed. These results confirm previous findings from Fiorillo et al. (2003) regarding the role of dopamine neurons in risk processing and previous findings from, e.g., Schultz (1998) regarding the role of dopamine neurons in RPE processing, but not regarding a possible role of these nuclei also in EV processing. Only the SN was additionally associated with outcome surprise, similarly to Matsumoto and Hikosaka (2009). As a control analysis (see Table S2), we also fit a GLM using the design of Preuschoff et al. (2006). In particular, we fit separate regressors for the first and second epoch after presenting the first number (where the first epoch lasted 1 second and the second epoch lasted 3 seconds). In these analyses, we found significant correlation with risk (in both epochs) and RPE across both the SN and the VTA. However, contrary to the results of our primary analysis, we also found significant correlation with EV in the second epoch with right-SN and left-VTA and no significant correlation with surprise. Note, however, that the high correlation between regressors in the first and second epochs (see Figure S5) might limit the sensitivity of our analysis given our particular task.

**Table 2.**
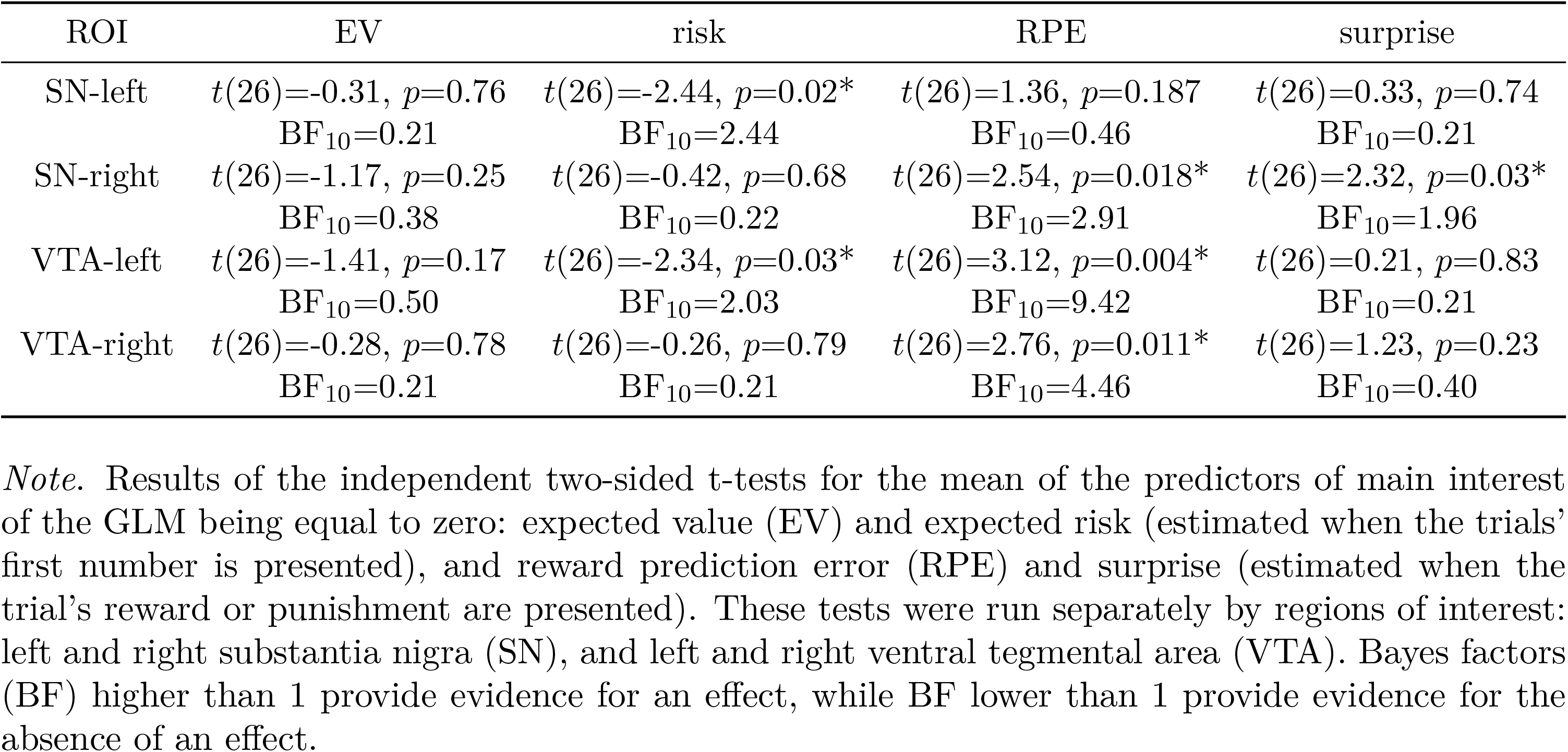
ROI-wise GLM results.

| ROI | EV | risk | RPE | surprise |
| --- | --- | --- | --- | --- |
| SN-left | $t(26)=-0.31, p=0.76$<br>BF<sub>10</sub>=0.21 | $t(26)=-2.44, p=0.02^*$<br>BF<sub>10</sub>=2.44 | $t(26)=1.36, p=0.187$<br>BF<sub>10</sub>=0.46 | $t(26)=0.33, p=0.74$<br>BF<sub>10</sub>=0.21 |
| SN-right | $t(26)=-1.17, p=0.25$<br>BF<sub>10</sub>=0.38 | $t(26)=-0.42, p=0.68$<br>BF<sub>10</sub>=0.22 | $t(26)=2.54, p=0.018^*$<br>BF<sub>10</sub>=2.91 | $t(26)=2.32, p=0.03^*$<br>BF<sub>10</sub>=1.96 |
| VTA-left | $t(26)=-1.41, p=0.17$<br>BF<sub>10</sub>=0.50 | $t(26)=-2.34, p=0.03^*$<br>BF<sub>10</sub>=2.03 | $t(26)=3.12, p=0.004^*$<br>BF<sub>10</sub>=9.42 | $t(26)=0.21, p=0.83$<br>BF<sub>10</sub>=0.21 |
| VTA-right | $t(26)=-0.28, p=0.78$<br>BF<sub>10</sub>=0.21 | $t(26)=-0.26, p=0.79$<br>BF<sub>10</sub>=0.21 | $t(26)=2.76, p=0.011^*$<br>BF<sub>10</sub>=4.46 | $t(26)=1.23, p=0.23$<br>BF<sub>10</sub>=0.40 |

*Note.* Results of the independent two-sided t-tests for the mean of the predictors of main interest of the GLM being equal to zero: expected value (EV) and expected risk (estimated when the trials' first number is presented), and reward prediction error (RPE) and surprise (estimated when the trial's reward or punishment are presented). These tests were run separately by regions of interest: left and right substantia nigra (SN), and left and right ventral tegmental area (VTA). Bayes factors (BF) higher than 1 provide evidence for an effect, while BF lower than 1 provide evidence for the absence of an effect.

**Figure 4.**
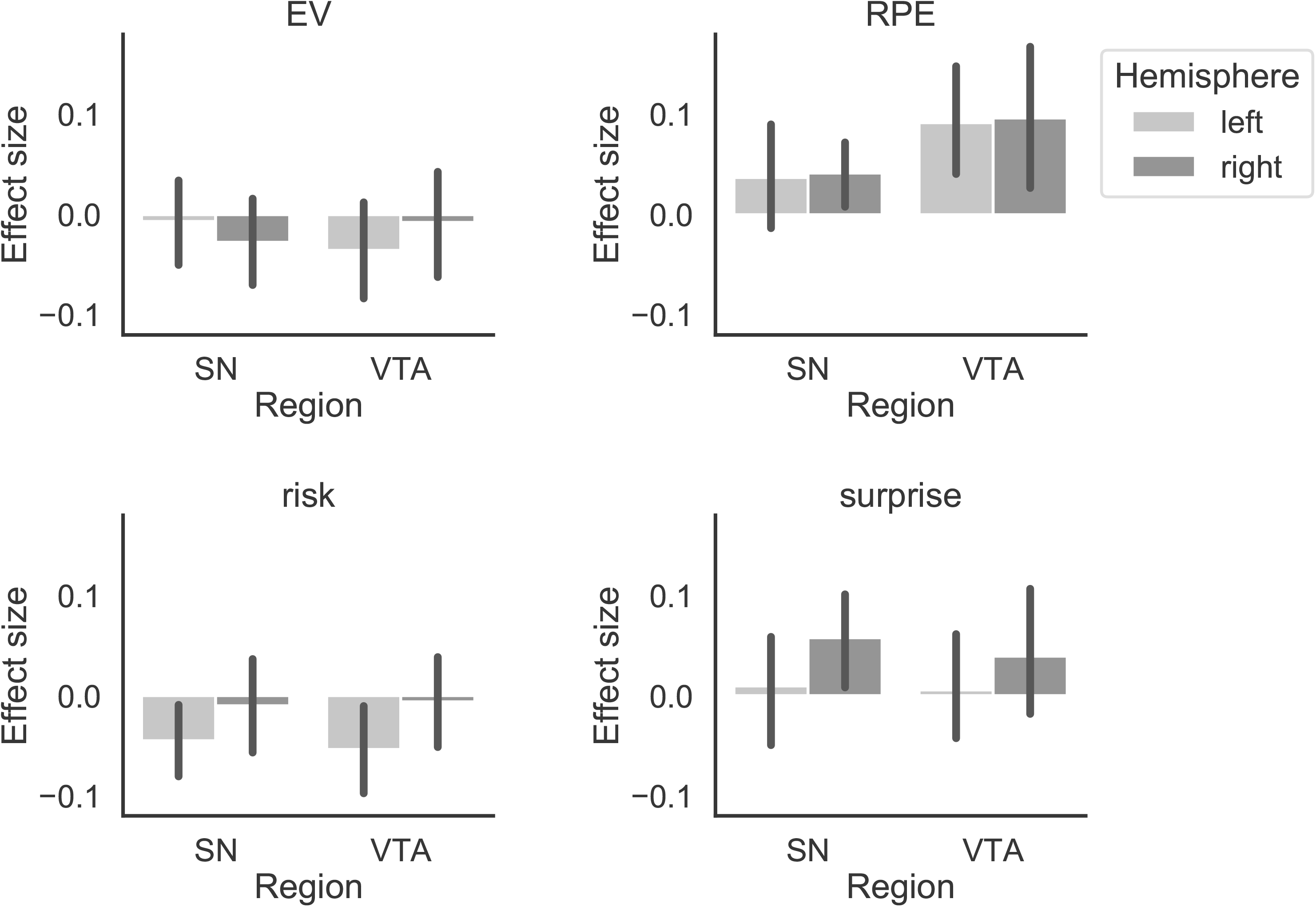
Average effect size across participants of the GLM on the time-series data extracted from the regions of interest (ROI): left and right substantia nigra (SN) and left and right ventral tegmental area (VTA). Different plots represent the predictors of main interest: expected value (EV) and expected risk (estimated when the trials’ first number is presented), and reward prediction error (RPE) and surprise (estimated when the trial’s reward or punishment are presented). Error bars represent 95% confidence intervals.

### Voxel-wise GLM

To explore other sub-cortical and cortical correlates of expectation- and feedback-related processes, we fit the same GLM on the whole-brain level. The results are shown in Table 3 and Figure 5 (see also Table S1 for automatic labeling based on cluster peak coordinates). After cluster correction, we found positive correlations with EV in the ventromedial prefrontal cortex, frontal pole, ventral striatum, and precuneous cortex, and negative correlations with EV in the thalamus. We found positive correlations with risk in the middle temporal gyrus and posterior insula, and negative correlations with risk in orbital frontal cortex, frontal lobe, and anterior insula. We found positive correlations with RPE in ventral striatum, orbital frontal cortex, midbrain, precuneus and anterior insula, and no negative correlations with RPE. Finally, we found positive correlations with surprise in the orbital frontal cortex, inferior frontal gyrus, superior temporal gyrus, and middle temporal gyrus, and negative correlations with surprise in precuneus and posterior insula. Even though we could not test for temporal differentiation in the anticipatory period (due to identifiability issues, see above), we could observe a spatial differentiation between EV and risk, confirming parts of the results from Preuschoff et al. (2006). We also observed a spatial differentiation between RPE and surprise.

**Table 3.** Results of the voxel-wise GLM after cluster-wise thresholding.

| Predictor | Cluster Index | Voxels | p | -log10(p) | Max/Min | Max/Min x (vox) | Max/Min y (vox) | Max/Min z (vox) |
| --- | --- | --- | --- | --- | --- | --- | --- | --- |
| EV (positive) | 5 | 871 | <0.001 | 8.57 | 3.56 | 12.0 | -55.5 | 25.5 |
| EV (positive) | 4 | 868 | <0.001 | 8.54 | 3.88 | -6.0 | 42.0 | -15.0 |
| EV (positive) | 3 | 283 | 0.002 | 2.60 | 3.56 | -7.5 | 16.5 | -6.0 |
| EV (positive) | 2 | 275 | 0.003 | 2.50 | 3.53 | 9.0 | 16.5 | -6.0 |
| EV (positive) | 1 | 237 | 0.01 | 2.01 | 3.78 | -31.5 | 51.0 | -16.5 |
| EV (negative) | 3 | 1274 | <0.001 | 11.80 | -3.67 | 6.0 | -27.0 | -1.5 |
| EV (negative) | 2 | 226 | 0.014 | 1.86 | -3.02 | -15.0 | -75.0 | -10.5 |
| EV (negative) | 1 | 218 | 0.018 | 1.75 | -3.01 | -18.0 | -76.5 | 3.0 |
| risk (positive) | 5 | 7344 | <0.001 | 31.00 | 4.67 | -19.5 | -45.0 | 18.0 |
| risk (positive) | 4 | 1054 | <0.001 | 6.62 | 3.82 | 49.5 | -25.5 | 15.0 |
| risk (positive) | 3 | 591 | <0.001 | 3.73 | 3.73 | 16.5 | 34.5 | -3.0 |
| risk (positive) | 2 | 365 | 0.01 | 2.01 | 4.05 | -19.5 | 33.0 | -4.5 |
| risk (positive) | 1 | 305 | 0.031 | 1.50 | 4.23 | 52.5 | -60.0 | -1.5 |
| risk (negative) | 7 | 2664 | <0.001 | 14.50 | -4.67 | 30.0 | 24.0 | -9.0 |
| risk (negative) | 6 | 1300 | <0.001 | 8.05 | -3.83 | 15.0 | 64.5 | -4.5 |
| risk (negative) | 5 | 1209 | <0.001 | 7.55 | -4.08 | 6.0 | -91.5 | 7.5 |
| risk (negative) | 4 | 950 | <0.001 | 6.05 | -3.75 | -31.5 | 16.5 | -18.0 |
| risk (negative) | 3 | 515 | 0.001 | 3.18 | -4.31 | -16.5 | 66.0 | -3.0 |
| risk (negative) | 2 | 467 | 0.002 | 2.82 | -3.93 | 67.5 | -30.0 | -6.0 |
| risk (negative) | 1 | 397 | 0.005 | 2.27 | -3.31 | -10.5 | 39.0 | 6.0 |
| RPE (positive) | 8 | 4234 | <0.001 | 24.10 | 4.44 | 12.0 | 16.5 | -4.5 |
| RPE (positive) | 7 | 2724 | <0.001 | 17.30 | 4.38 | -10.5 | 18.0 | -12.0 |
| RPE (positive) | 6 | 526 | <0.001 | 4.04 | 3.70 | -27.0 | -25.5 | 13.5 |
| RPE (positive) | 5 | 506 | <0.001 | 3.87 | 3.59 | 40.5 | 55.5 | -10.5 |
| RPE (positive) | 4 | 398 | 0.001 | 2.91 | 4.30 | -42.0 | 48.0 | -15.0 |
| RPE (positive) | 3 | 390 | 0.001 | 2.84 | 3.64 | 12.0 | -25.5 | -10.5 |
| RPE (positive) | 2 | 314 | 0.008 | 2.11 | 3.96 | 63.0 | -25.5 | 25.5 |
| RPE (positive) | 1 | 273 | 0.02 | 1.69 | 3.81 | 16.5 | -54.0 | 22.5 |
| surprise (positive) | 4 | 1026 | <0.001 | 9.89 | 4.33 | 57.0 | 22.5 | 6.0 |
| surprise (positive) | 3 | 871 | <0.001 | 8.58 | 4.41 | 51.0 | -33.0 | 0.0 |
| surprise (positive) | 2 | 464 | <0.001 | 4.70 | 3.73 | -58.5 | -25.5 | -6.0 |
| surprise (positive) | 1 | 328 | 0.001 | 3.16 | 3.54 | 54.0 | 13.5 | -21.0 |
| surprise (negative) | 3 | 1291 | <0.001 | 12.00 | -4.04 | -22.5 | -54.0 | 7.5 |
| surprise (negative) | 2 | 635 | <0.001 | 6.45 | -4.16 | 10.5 | -51.0 | 13.5 |
| surprise (negative) | 1 | 250 | 0.007 | 2.18 | -3.31 | 48.0 | 0.0 | 12.0 |

*Note.* Clusters surviving thresholding. We report the number of voxels, cluster probability, log probability, activation and MNI coordinate of the activation peak voxel in a cluster.

**Figure 5.**
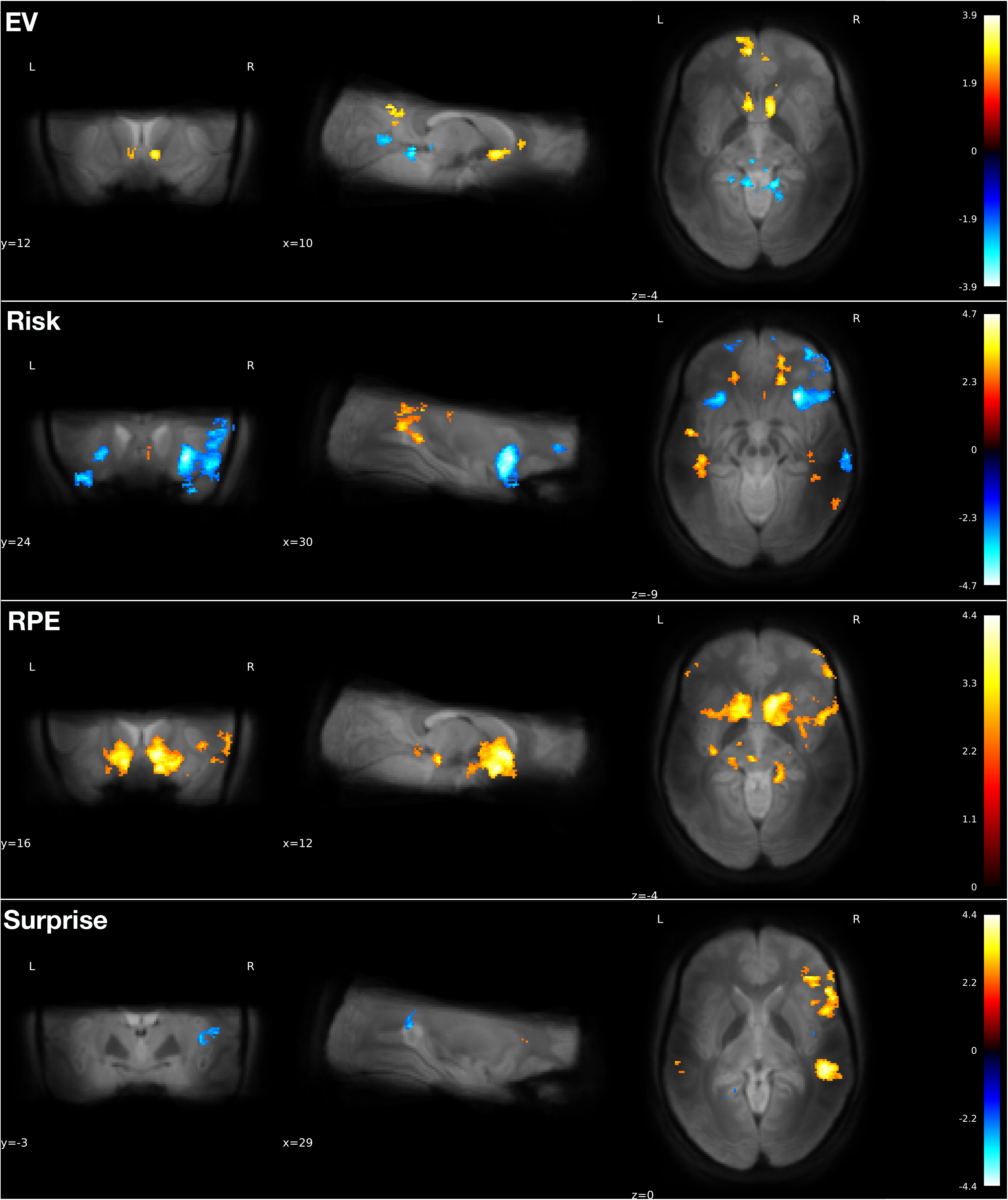
Results of the voxel-wise GLM after cluster correction, and overlapped onto the mean functional image across participants and volumes. Each row corresponds to the predictors of main interest: expected value (EV) and expected risk (estimated when the trials’ first number is presented), and reward prediction error (RPE) and surprise (estimated when the trial’s reward or punishment are presented).

## Discussion

Understanding the dopamine circuit is of great importance for both clinical and cognitive neuroscience. First of all, the loss of dopaminergic neurons is associated with Parkinson’s disease symptoms (Fearnley & Lees, 1991; Frank, 2006a) and dysregulations in the human dopamine circuit are known to play a role in drug addiction (Everitt & Robbins, 2005) and pathological gambling (Bergh, Eklund, Södersten, & Nordin, 1997). Moreover, the dopamine signal reflects different aspects of rewards, including the anticipation of risk and the mismatch between predictions and outcomes (Schultz, 2015). While dopamine neurons are situated mostly in the midbrain, they are part of a much greater and complex circuit, involving different cortical and subcortical areas (Frank, 2006b; Haber & Knutson, 2010; Watabe-Uchida, Eshel, & Uchida, 2017). By transmitting information about changes in reward expectations and risk in the environment to areas important for action execution and learning, dopamine likely plays a crucial role in adaptive behavior, that is, for survival in a dynamic environment, with limited resources and obstacles to avoid.

To date, most human studies have focused on the target areas (both cortical and subcortical) of the dopamine neurons because of methodological challenges. An exception was the study of Zaghloul et al. (2009): Using microelectrode recordings during deep brain stimulation surgery in Parkinson’s disease patients, they found SN activation in line with the RPE. Importantly, human studies that investigated the activity of dopamine nuclei using fMRI provided incomplete and partially contradicting results. In this paper, we presented the results of a 7T fMRI study involving human participants performing a number-guessing task. To the best of our knowledge, this was the first study to investigate the functional role of both the VTA and the SN using UHF-MRI to acquire high-quality, high-resolution functional and structural images. While previous studies in these areas focused on expected gains or losses and on the RPE signals, we extended the analysis to expected risk and to surprise. This was based on previous electrophysiological and fMRI studies that either found this signal in the VTA/SN or in their target areas (e.g., Fiorillo et al., 2003; Hayden, Heilbronner, Pearson, & Platt, 2011; Preuschoff et al., 2006). While we found no evidence for a linear correlation between reward anticipation (involving both gains and losses) and VTA or SN activation, we did find evidence for a RPE signal in both regions, as well as for expected risk signal. Similarly to Matsumoto and Hikosaka (2009), who found a functional dissociation of VTA and SN, we also found a surprise signal in the SN but not in the VTA.

Given previous findings (Fiorillo et al., 2003) and theoretical considerations (as a reward predicting cue could elicit already a RPE, when the reward expectations through the whole experiment are known; see Hare, O’Doherty, Camerer, Schultz, & Rangel, 2008), one might expect to find EV signals in the SN/VTA. Since participants were explicitly instructed that the initial bet’s outcome was random, there was perhaps less focus on the action and more on the reward structure of the task (i.e., the distribution of outcome one can expect given a certain number and choice pair). Note, however, that we did find positive correlations with EV in the ventromedial prefrontal cortex and ventral striatum, in line with previous studies inspecting value signaling in the cortex (Bartra et al., 2013; Schoenbaum, Takahashi, Liu, & McDannald, 2011).

The presence of a full RPE signal in both the VTA and the SN confirms previous results in animal studies (Schultz, 2015), although most of them are based on signal from the lateral part of the the VTA alone (Eshel, Tian, Bukwich, & Uchida, 2016). It also clarifies previous results on the VTA/SN signals in fMRI human studies (D’Ardenne et al., 2008; Pauli et al., 2015; Zhang et al., 2017). For instance, D’Ardenne et al. (2008) only found evidence for a positive RPE in VTA and not in SN. We also found an RPE signal in ventral striatum, orbital frontal cortex, and anterior insula, confirming previous fMRI results that looked at dopamine target areas (Bartra et al., 2013).

Here, we showed the presence of a risk signal in both the VTA and the SN, in line with electrophysiological studies in non-human animals (Fiorillo et al., 2003). We also found a risk signal in insula and orbital frontal cortex, confirming previous fMRI studies linking these areas to the coding of risk (Brown & Braver, 2018; Preuschoff et al., 2006).

The presence of a surprise signal in the SN and not in the VTA fits remarkably well with results from the animal literature (Matsumoto & Hikosaka, 2009) and with the framework proposed by Bromberg-Martin, Matsumoto, and Hikosaka (2010). In this framework, there are two distinct functional groups of dopamine neurons, a motivational value group, that shows the standard RPE response, and a motivational salience group, that reflects how unexpected outcomes are – positive or negative alike. Cells of the first group are situated more in the ventromedial part of the SNc and throughout the VTA, while cells of the second group are situated more in the dorsolateral part of the SNc as well as in the medial VTA. While SNc cells project more to sensorimotor dorsolateral striatum, VTA cells project more to ventral striatum. Beyond our ROIs, we also found correlations between surprise and posterior (but not anterior) insula.

Both the SN and the VTA are relatively small subcortical structures (around 511 mm^3^ and 138 mm^3^, respectively, see Table 1), they are adjacent to each other as well as to other nuclei with related functions, such as the red nucleus and the subthalamic nucleus, and they are susceptible to other possible sources of noise, such as the physiological noise in the cerebrospinal fluid. The small dimension of the nuclei and their spatial contiguity increase the risk of confusing the signal from different regions (de Hollander et al., 2015; Trutti et al., 2019). To be able to more reliably extract and separate the signals from the VTA and the SN, we therefore drew individual masks, based on 0.7 mm isotropic, multimodal, anatomical images that were acquired for each participant in a separate session. By restricting the analyses to the individual space, we also prevented misalignment issues that usually occur when transforming individual images to a group or standard space. To define the final masks, we adopted a rather conservative approach, by keeping the intersection of the masks drawn by two independent and trained raters. To illustrate the importance of these precautions, we compared our masks to previously proposed VTA and SN probabilistic masks in the standard space. In particular, we considered the SN subdivisions proposed by Zhang et al. (2017) and the VTA and the SN subdivisions proposed by Pauli et al. (2018). We found that when transforming these masks to the individual space – as it is usually done during ROI signal extraction – the signal from the VTA and the SN is indeed partially mixed. This can have serious impact on the interpretation of the results of an fMRI study. For instance, Zhang et al. (2017) reported an RPE signal in the medial part of the SN, which – according to our analyses and results – is the part that overlaps the most with the VTA, and a surprise signal in the lateral part of the SN. To be able to draw strong conclusions on the functional specificity of – in this case – SN subdivisions, we would thus argue that it is preferable to have individually drawn masks.

Future studies could attempt to distinguish between the pars compacta and reticulata of the SN, as dopamine neurons are mainly situated in the pars compacta (Roeper, 2013). However, these two parts are virtually indistinguishable based on MRI contrast alone (see Figure 2). Therefore, to avoid making an arbitrary decisions on where to set a border between the two, we considered the SN as one structure. By combining different methodologies (i.e., diffusion MRI) future studies might be able to shed light on SN functional subdivisions.

Another limitation of the present study relies in the nature of the BOLD signal. Since the BOLD response measured in fMRI is an indirect measure of neuronal activity and is mainly thought to measure signals input and local processing of neurons rather than their output (Logothetis & Wandell, 2004), it is important to integrate results from different methodologies and species in order to understand the complexity of the dopaminergic circuit as a whole.

In sum, in this study we used novel methodologies to investigate how the brain processes gains and losses and updates expectations based on experience. We were able to show a risk signal in the dopamine nuclei and provided evidence for a full RPE signal in the presence of both gains and losses, thus clarifying previous results of human fMRI studies. This study opens the way to a better understanding of the dopamine circuit in the human brain, especially regarding the functional specificity of the SN and the VTA (or of their subregions) in reward-based decision making and adaptive behavior.

## Materials & Methods

### Participants and procedure

Twenty-seven participants [8 male (mean age=24.7, SD=5.0, min=19, max=35), 19 female (mean age=24.4, SD=4.7, min=19, max=35)] took part in the experiment. The study was approved by the ethics committee of the University of Amsterdam. All participants completed two separate sessions, one to obtain multimodal, 0.7 mm isotropic structural data, and one to obtain 1.5 mm isotropic functional data while participants engaged in a number-guessing task. All participants were recruited from the University of Amsterdam subject pool, via flyers and posters at the Spinoza center for Neuroimaging and at the Academic Medical Center in Amsterdam, and via advertisements in the magazine of the Dutch Parkinson Society. All participants were required to be MRI compatible, between 18 and 40 years old, right-handed, without previous history of psychiatric conditions or neurological diseases, and to have normal or corrected-to-normal vision. Before taking part in the sessions they gave written consent, and, before the second session, they received written instructions for the behavioral task. Before going in the MRI scanner, they all completed a training session in which they could try the experiment on a computer, and were given a written questionnaire to test their comprehension of the probability of winning and losing in each scenario of the behavioral task. All participants were given 20 euros as compensation for the second session. In the second session they could win or lose up to 7 euros based on their performance in the task, which were either added or subtracted from an additional endowment of 10 euro.

### Data acquisition

All images were acquired on a Philips Achieva 7T MRI scanner, situated at the Spinoza Centre for Neuroimaging in Amsterdam (Netherlands), using a Nova Medical 32-channel head array coil. During the first session, participants could choose whether to watch a movie or not. During the second session, the number-guessing task was presented using PsychoPy (Peirce, 2007).

#### Structural MRI

T_1_-weighted, 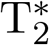-weighted, and Quantitative Susceptibility Mapping (QSM, Langkammer et al., 2012) images were simultaneously obtained using a multi-echo magnetization-prepared rapid gradient echo (ME-MP2RAGE) sequence (Caan et al., 2018; Metere, Kober, Möller, & Schäfer, 2017). The sequence parameters were: T_*I*,1_ = 670 ms, T_*I*,2_ = 3675.4 ms, T_*R*,1_ = 6.2 ms, T_*R*,2_ = 31 ms, T_*E*,1_ = 3 ms, T_*E*,2_ = [3, 11.5, 19, 28.5 ms], T_*R,MP*_ 2_*RAGE*_ = 6778 ms, flip angle_1_: 4°, flip angle_2_: 4^*◦*^, bandwidth: 404.9 MHz, acceleration factor SENSE: 2, FOV = 205 × 205 × 164 mm^3^, acquired voxel size: .7 × .7 × .7 mm^3^, acquisition matrix: 292 × 290, reconstructed voxel size: .64 × .64 × .70 mm^3^, turbo factor: 150 (resulting in 176 shots). The total acquisition time was 19.53 min.

#### Functional MRI

The functional MRI protocol was an adaptation of Protocol 3 as reported by (de Hollander et al., 2017), originally designed for a 7T Siemens scanner located at the Max Planck Institute for Human Cognitive and Behavioral Sciences in Leipzig, Germany. This protocol was used to optimize the tSNR in iron-rich nuclei in the human midbrain. The present protocol consisted of 2 runs of 719 volumes with 30 slices. The acquisition time was 23.97 min per run. Other parameters were T_*R*_ = 2,000 ms, T_*E*_ = 17 ms, flip angle: 60^*◦*^, bandwidth: 2226.2 Hz, voxel size: 1.5 × 1.5 × 1.5 mm ^3^, FOV = 192 × 192 × 49 mm^3^, SENSE acceleration factor, P-reduction (AP): 3, matrix size: 128 × 128. To acquire images with such TE, TR, and voxel-size, the protocol did not employ Fat suppression, and, to increase SNR, the protocol did not employ Partial Fourier. After the first run, an EPI image with opposite phase coding direction as compared to the functional scan was acquired to help correcting for geometric distortions due to inhomogeneities in the B0 field using the TOPUP technique during preprocessing (see below).

### Number-guessing task

The number-guessing task used in the present study is an adaptation of the task by Preuschoff et al. (2006). In each trial (Figure 1A), two numbers were sampled one after the other from the set 1, 2, 3, 4, 5 without replacement. At the beginning of each trial, before seeing both numbers, participants were asked to bet which of the two numbers will be higher: They could win 5 euro if their bet (i.e., their prediction) was correct, and lose 5 euro otherwise. Participants were also instructed that the sampling was (pseudo-) random and that their choice could not influence sampling. The texts *“Second number is HIGHER*.” and *“Second number is LOWER*.” appeared on the left and right side of the screen, respectively (the position was counterbalanced across participants), and participants had to press either a left or a right button to place their bet. They could do so within 1 second, otherwise a bet would be placed for them at random. The choice (either the participant’s or the random one) was then indicated by presenting a black frame around the corresponding text for another second.

The first number was subsequently shown for 2 seconds. Based on this first number, participants can update the probability to win or lose (both 50% at the beginning of the trial). For example, if a bet is placed on the second number being higher than the first number, and the first number is revealed to be 2, then three out of the four remaining numbers (i.e., 3, 4, and 5) lead to winning (*p*_*winning*_ = 75%), while only one number (i.e., 1) leads to losing (*p*_*losing*_ = 25%). The expected value (EV) of the gamble is calculated as:

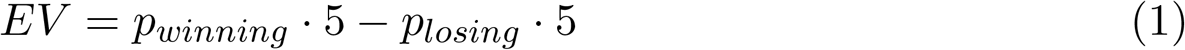

and in this case is thus 5⋅0.75−5⋅0.25 = 2.5 euros. The risk, often defined as the variance of the possible outcomes (Markowitz, 1952), is thus 4.3. Note that, when the first number is 3, the probability to win remains 50%, the EV remains 0, and the risk is highest, equal to 5. On the contrary, when the first number is either 1 or 5, participants already know whether they will lose or win (depending on what the bet was), therefore the EV is either −5 or 5 euros and the risk is always 0. Since we were interested in neural correlates of both EV and risk, it is a crucial aspect of this design that EV and risk are not correlated (Figure 1B).

At last, the second number is shown for 2 seconds, together with the corresponding gain or loss. At this point, the reward prediction error (RPE) is calculated:

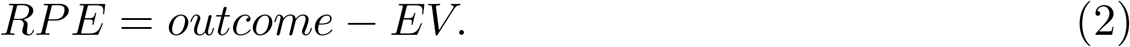

In the example above (i.e., bet on 2nd number being higher; first number is 2), if the second number is 3, the reward is 5 euros and the reward prediction error is 5 − 2.5 = 2.5 euros. The surprise, defined as the absolute value of the reward prediction error (i.e., the reward expectation after the first number) as in Schultz (2015) and in Hayden et al. (2011), is thus |5 − 2.5| = 2.5. Since we were also interested in neural correlates of both RPE and surprise, it was also crucial that they were uncorrelated. This was the case, since RPE ranged between −7.5 and 7.5 and its distribution over trials was symmetrically centered around 0, and surprise was simply its absolute value.

The experiment consisted of 120 trials, divided in two blocks. In each block, 5 test trials were included to encourage participants to remain attentive throughout the experiment. In these trials, instead of showing the reward, we asked participants to indicate whether they won or lost. To correctly respond to this question, they needed to remember both their bet and the first number. At the end of the experiment, we randomly selected one of the 110 regular trials, and participants received the corresponding reward (i.e., 5 or −5 euros), plus 2 additional euros if they responded correctly to at least 8 of the 10 test trials, otherwise we subtracted 2 euros to the final reward. Between each event in each trial, and at the beginning of each trial, a fixation cross was presented for a period of time between 4 and 10 seconds, drawn from a truncated exponential distribution. The long inter-stimuli intervals were crucial to allow separating the BOLD signals associated with the first and the second numbers (i.e., signals related to either expectations or feedback processing).

### Behavioral analysis

Because choices are not influencing the chance of winning or losing in this task, behavioral analyses had the purpose to check the quality of the data for the fMRI analyses. The most important indicator of data quality was the accuracy in the test trials: Blocks in which participants made more than two out of five mistakes were discarded, where misses also counted as mistakes. Another important indicator was the number of missed bets: Blocks in which participants missed more than ten out of 60 bets were discarded. Finally, we checked the percentage of right vs. left responses. Because the position of the texts corresponding to the specific bets was counterbalanced across – but fixed within – participants, a similar number of right and left responses needed to be made for a balanced design. Blocks in which participants made less than ten right or more than fifty right (out of 60) choices were discarded.

### Structural and functional MRI data preprocessing

Registration and preprocessing were performed using FMRIPREP version 1.0.6 (Esteban et al., 2018), a Nipype (Gorgolewski et al., 2011) based tool. Registration across session was done by registering the functional images (from the second session) to the T_1_-weighted structural image multiplied by the first echo of the 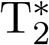-weighted structural images (from the first session). Because the T_1_-weighted, 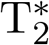-weighted, and QSM structural images were acquired simultaneously during the same scan in the first session, there was no need to co-register them first.

Structural images were corrected for intensity non-uniformity using N4 Bias Field Correction (Tustison et al., 2010) and skull-stripped using antsBrainExtraction.sh. Spatial normalization to the ICBM 152 Nonlinear Asymmetrical template (Fonov, Evans, McKinstry, Almli, & Collins, 2009) was performed through nonlinear registration with the antsRegistration tool of ANTs v2.1.0 (Avants, Epstein, Grossman, & Gee, 2008), using brain-extracted versions of both T_1_-weighted volume and template. Brain tissue segmentation of cerebrospinal fluid (CSF), white-matter (WM) and gray-matter (GM) was performed on the brain-extracted T_1_-weighted image using *fast* (FSL v5.0.9) (Zhang, Larcher, Misic, & Dagher, 2001). Functional data was motion corrected using mcflirt (FSL v5.0.9, Jenkinson, Bannister, Brady, & Smith, 2002). Distortion correction was performed using an implementation of the TOPUP technique (Andersson, Skare, & Ashburner, 2003) using 3dQwarp (AFNI v16.2.07, Cox, 1996). This was followed by co-registration to the corresponding T_1_-weighted image using boundary-based registration Greve and Fischl (2009) with 9 degrees of freedom, using flirt (FSL). Motion correcting transformations, field distortion correcting warp, BOLD-to-T_1_-weighted transformation and T_1_-weighted-to-template (MNI) warp were concatenated and applied in a single step using antsApplyTransforms (ANTs v2.1.0) using Lanczos interpolation.

Physiological noise regressors were extracted applying CompCor (Behzadi, Restom, Liau, & T.Liu, 2007). Principal components were estimated for the anatomical CompCor (aCompCor). A mask to exclude signal with cortical origin was obtained by eroding the brain mask, ensuring it only contained subcortical structures. Six tCompCor components were then calculated including only the top 5% variable voxels within that subcortical mask. For aCompCor, six components were calculated within the intersection of the subcortical mask and the union of CSF and WM masks calculated in T_1_-weighted space, after their projection to the native space of each functional run. FD was calculated for each functional run using the implementation of Nipype.

The preprocessing and registration output was visually inspected for each subject using the html output files of FMRIPREP. Functional data quality was assessed using MRIQC (Esteban et al., 2017) prior preprocessing, to check for visual artifacts and excessive head movements. Finally, after preprocessing and registration, tSNR maps were computed using Nipype to assess the tSNR across the ROIs.

### Anatomical segmentation

One main aim of the present study was to obtain anatomically precise masks in the individual space for the two ROIs: the ventral tegmental area (VTA) and the substantia nigra (SN). Because of its relatively high iron concentration, the SN is most discernible in QSM images (Keuken et al., 2014), as shown in the first row of Figure 2. Unlike the SN, the VTA lacks clear anatomical borders (Trutti et al., 2019). Segmentation can be performed, however, by exclusion from the neighboring iron-rich nuclei (i.e., the SN and the red nucleus, RN) and the CSF, so both should be clearly visible. The CSF is not visible in the QSM image. It is, however, clearly visible in the T_1_-weighted image (see Figure 2, third row). To ease and improve the segmentation process, we therefore combined the 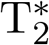-weighted and T_1_-weighted images, by first normalizing them within the midbrain area (i.e., a pre-selected area of 1.6 × 1.6 × 3.08 cm^3^) and finally summing them up. The result can be seen in the bottom row of Figure 2: The QSM images in the first row show a high contrast for iron rich areas, such as the SN, the red nucleus (situated above and posterior to the SN), and the subthalamic nucleus (situated above and anterior to the SN); the 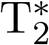-weighted and T_1_-weighted images (second and third row) highlight, respectively, iron rich areas and the CSF; their sum (fourth row) thus allows to segment the VTA, as it is mainly defined by the border it shares with these regions (which are hard to visualize within the same contrast).

Manual segmentation was performed using FSLView version 3.0.2, by two independent and trained researchers (one of which is the first author of this study). Only the voxels that were marked by both researchers were kept in the final masks, that is, the conjunction masks. To assess inter-rater reliability (i.e., the agreement between the two researcher), we computed the Dice score (Dice, 1945) separately for each participant, hemisphere, and structure. The Dice score is computed as the ratio between the union of the two areas and the conjunction of the two areas. It therefore depends on the average dimension of the structure (with smaller structures usually having lower scores) and has to be interpreted accordingly. Scores approaching 1 indicate perfect agreement between raters, while scores close to 0 indicate no agreement between raters.

Drawing individual masks for each subject and area is a time- and resource-consuming process: High resolution structural images need to be acquired first, and then two trained researchers need to complete a lengthy segmentation process. To forgo this costly approach, SN (Keuken et al., 2014) and VTA (Pauli et al., 2018) MRI atlases have been published in recent years. These atlases consists of probabilistic maps of different ROIs in MNI space, and can be thus transformed in the individual space to extract the signal from these regions. The disadvantage of this less resource-intensive approach, however, is a potential loss of sensitivity and specificity due to misalignment between the individual and the standard spaces, as well as individual differences. To quantify the loss of information in this process, we transformed the three SN subregions proposed by Zhang et al. (2017), based on the 33% thresholded probabilistic masks proposed by Keuken et al. (2014), to the individual space and measured the overlap with our individual VTA masks as the number of voxels in common, divided by the overall area. A similar procedure was done with the proposed VTA and SN subdivisions of Pauli et al. (2018), using their deterministic atlas (50% thresholded).

### fMRI data analysis

We extracted the fMRI signal for each time point within the ROIs (i.e., left and right SN and VTA) for each subject and computed its average time course for each ROI separately. We then fitted a GLM to the resulting time series for every region, participant, and block using statsmodels (Seabold & Perktold, 2010). Specifically, we used the GLSAR AR(1) model, to account for autocorrelation. The design matrices were constructed using Nistats (https://nistats.github.io/index.html). In the design matrices, the following events were convolved with the canonical, double-gamma hemodynamic response function (HRF): the bet at the beginning of the trial, the appearance of the first number, the appearance of the second number in regular trials, and the appearance of the second number in test trials. On top of these, we added four parametric regressors: EV and risk (with onsets at the appearance of the first number and as amplitude the normalized EV and risk of each trial), and RPE and surprise (with onsets at the appearance of the second number and as amplitude the normalized RPE and surprise of each trial). The duration of the parametric regressors, together with their intercepts (i.e., the appearance of the first and second number), was set to 2 seconds, as this was the time of presentation of the numbers on the screen. Additional nuisance parameters were the six aCompCor, FD, six head movement variables provided by *fmriprep*, and cosine regressors for high-pass temporal filtering. No spatial smoothing was used. After averaging across blocks, we performed independent two-sided t-tests, separately by ROIs and hemisphere (i.e., left vs. right) for the mean of the parameters corresponding to EV, risk, RPE, and surprise being equal to zero. We also estimated the equivalent Bayesian t-tests, as implemented in the BayesFactor R library (https://cran.r-project.org/web/packages/BayesFactor/index.html), as it allows quantifying evidence in favor of the null hypothesis and therefore complements the frequentist analyses.

For the exploratory and control analyses, we estimated the same GLMs as on the ROIs, using a mass-univariate, voxel-wise approach with Nistats (https://nistats.github.io/index.html). At the level of individual runs, we used a smoothing Gaussian kernel with a FWHM of 3.0 mm. At the participant level, we estimated the size of the baseline contrasts of the parameter estimates of EV, risk, RPE, and surprise. These participant-wise contrasts of parameter estimates (COPE) were then transformed to the MNI space and used in the third and final group-level analysis. Finally, we performed a Gaussian Random Field cluster analysis on the resulting four z-maps (EV, risk, RPE, and surprise), using FSL *cluster tool*. For these analyses, we set an input threshold of 2.3 and a cluster-wise threshold of p*<*.05.

## Acknowledgements

We would like to thank Josephine Groot and Martijn Mulder for helping with collecting the data and recruiting participants, Wietske van der Zwaag for helping setting up the fMRI protocol, and Gilles de Hollander for consultation concerning fMRI in the midbrain. Further thanks go to Anne Trutti, for sharing her knowledge of the VTA anatomy and for helping with VTA segmentation, and to Anneke Alkemade, for helping with the SN segmentation.

## Footnotes

1 Defined as the ratio between the number of voxels in common and the number of voxels in the subdivision.

## Notes

This research is supported by the Swiss National Science Foundation (mobility grant number P1BSP1_172017), the Department of Psychology of the University of Amsterdam, the Netherlands Organisation for Scientific Research (project number 14017), and an ERC grant from the European Research Council (B.U.F.).

https://osf.io/4vjta/

## References

Andersson, J. L. R., Skare, S., & Ashburner, J. (2003). Symmetric diffeomorphic image registration with cross-correlation: Evaluating automated labeling of elderly and neurodegenerative brain. NeuroImage, 20(2), 870–888. doi: 10.1016/S1053-8119(03)00336-7

Arias-Carrión, O., Stamelou, M., Murillo-Rodríguez, E., Menéndez-Gonzáles, M., & Pöppel, E. (2010). Dopaminergic reward system: A short integrative review. International Archives of Medicine, 3(24), 1–6. doi: 10.1186/1755-7682-3-24

Avants, B. B., Epstein, C. L., Grossman, M., & Gee, J. C. (2008). Symmetric diffeomorphic image registration with cross-correlation: Evaluating automated labeling of elderly and neurodegenerative brain. Medical Image Analysis, 12(1), 26–41. doi: 10.1016/j.media.2007.06.004

Bartra, O., McGuire, J. T., & Kable, J. W. (2013). The valuation system: A coordinate-based meta-analysis of bold fmri experiments examining neural correlates of subjective value. NeuroImage, 76, 412–427. doi: 10.1016/j.neuroimage.2013.02.063

Bayer, H. M., & Glimcher, P. W. (2005). Midbrain dopamine neurons encode a quantitative reward prediction error signal. Neuron, 47(1), 129–141.

Behzadi, Y., Restom, K., Liau, J., & T.Liu, T. (2007). A component based noise correction method (CompCor) for BOLD and perfusion based fMRI. NeuroImage, 37(1), 90–101. doi: 10.1016/j.neuroimage.2007.04.042

Bergh, C., Eklund, T., Södersten, P., & Nordin, C. (1997). Altered dopamine function in pathological gambling. Psychological Medicine, 27(2), 473–475. doi: 10.1017/S0033291796003789

Bromberg-Martin, E. S., Matsumoto, M., & Hikosaka, O. (2010). Dopamine in motivational control: Rewarding, aversive, and alerting. Neuron, 68(5), 815–834. doi: 10.1016/j.neuron.2010.11.022

Brown, J., & Braver, T. (2018). Risk prediction and aversion by anterior cingulate cortex Cognitive, Affective, & Behavioral Neuroscience, 7(4), 266–277. doi: 10.3758/CABN.7.4.266

Caan, M., Bazin, P.-L., Fracasso, A., Marques, J., Dumoulin, S., & van der Zwaag, W. (2018). MP2RAGEME: T1, T*_2_ and QSM mapping in one sequence at 7 Tesla. (Poster presented at the Joint Annual Meeting ISMRM-ESMRMB, Paris, France)

Clithero, J. A., & Rangel, A. (2014). Informatic parcellation of the network involved in the computation of subjective value. Social Cognitive and Affective Neuroscience, 9(9), 1289–1302. doi: 10.1093/scan/nst106

Cox, R. W. (1996). AFNI: Software for analysis and visualization of functional magnetic resonance neuroimages. Computers and Biomedical Research, 29(3), 162–173. doi: 10.1006/cbmr.1996.0014

D’Ardenne, K., McClure, S. M., Nystrom, L. E., & Cohen, J. D. (2008). BOLD responses reflecting dopaminergic signals in the human ventral tegmental area. Science, 319(5867), 1264–1267. doi: 10.1126/science.1150605

de Hollander, G., Keuken, M. C., & Forstmann, B. U. (2015). The subcortical cocktail problem; mixed signals from the subthalamic nucleus and substantia nigra. PloS One, 10(2), 1–18. doi: 10.1371/journal.pone.0120572

de Hollander, G., Keuken, M. C., van der Zwaag, W., Forstmann, B. U., & Trampel, R. (2017). Comparing functional MRI protocols for small, iron-rich basal ganglia nuclei such as the subthalamic nucleus at 7T and 3T. Human Brain Mapping, 38(6), 3226–3248. doi: 10.1002/hbm.23586

Dice, L. R. (1945). Measures of the amount of ecologic association between species. Ecology, 26(3), 297–302. doi: 10.2307/1932409

Drayer, B., Burger, P., Darwin, R., Riederer, S., Herfkens, R., & Johnson, G. (1986). Mri of brain iron. American Journal of Roentgenology, 147(1), 103–110.

Eapen, M., Zald, D. H., Gatenby, J. C., Ding, Z., & Gore, J. C. (2011). Using high-resolution MR imaging at 7T to evaluate the anatomy of the midbrain dopaminergic system. American Journal of Neuroradiology, 32(4), 688–694. doi: 10.3174/ajnr.A2355

Eshel, N., Tian, J., Bukwich, M., & Uchida, N. (2016). Dopamine neurons share common response function for reward prediction error. Nature Neuroscience, 19, 479–486. doi: 10.3174/ajnr.A2355

Esteban, O., Birman, D., Schaer, M., Koyejo, O. O., Poldrack, R. A., & Gorgolewski, K. J. (2017). MRIQC: Advancing the automatic prediction of image quality in MRI from unseen sites. PloS One, 12(9), 1–21. doi: 10.1371/journal.pone.0184661

Esteban, O., Markiewicz, C., Blair, R. W., Moodie, C., Isik, A. I., Aliaga, A. E., & Gorgolewski, K. J. (2018). FMRIPrep: a robust preprocessing pipeline for functional MRI. doi: 10.1101/306951

Everitt, B. J., & Robbins, T. W. (2005). Neural systems of reinforcement for drug addiction: From actions to habits to compulsion. Nature Neuroscience, 8(11), 1481–1489. doi: 10.1038/nn1579

Fearnley, J. M., & Lees, A. J. (1991). Ageing and Parkinson’s disease: Substantia nigra regional selectivity. Brain, 114(5), 2283–2301. doi: 10.1093/brain/114.5.2283

Fiorillo, C. D., Tobler, P. N., & Schultz, W. (2003). Discrete coding of reward probability and uncertainty by dopamine neurons. Science, 299(5614), 1898–1902. doi: 10.1126/science.1077349

Fonov, V. S., Evans, A. C., McKinstry, R. C., Almli, C. R., & Collins, D. L. (2009). Unbiased nonlinear average age-appropriate brain templates from birth to adulthood. NeuroImage, 4(Supplement 1), 39–41. doi: 10.1016/S1053-8119(09)70884-5

Forstmann, B. U., de Hollander, G., van Maanen, L., Alkemade, A., & Keuken, M. C. (2017). Towards a mechanistic understanding of the human subcortex. Nature Reviews Neuroscience, 18(1), 57–65.

Fouragnan, E., Retzler, C., & Philiastides, M. G. (2018). Separate neural representations of prediction error valence and surprise: Evidence from an fMRI meta-analysis. Human Brain Mapping, 39, 2887–2906. doi: 10.1002/hbm.24047

Frank, M. J. (2006a). Dynamic dopamine modulation in the basal ganglia: A neurocomputational account of cognitive deficits in medicated and nonmedicated Parkinsonism. Journal of Cognitive Neuroscience, 17(1), 17–52. doi: 10.1162/0898929052880093

Frank, M. J. (2006b). Hold your horses: a dynamic computational role for the subthalamic nucleus in decision making. Neural Networks, 19(8), 1120–1136.

Gorgolewski, K., Burns, C. D., Madison, C., Clark, D., Halchenko, Y. O., Waskom, M. L., & Ghosh, S. S. (2011). Nipype: A flexible, lightweight and extensible neuroimaging data processing framework in Python. Frontiers in Neuroinformatics, 5(13), 1–15. doi: 10.3389/fn-inf.2011.00013

Greve, D. N., & Fischl, B. (2009). Accurate and robust brain image alignment using boundary-based registration. NeuroImage, 48(1), 63–72. doi: 10.1016/j.neuroimage.2009.06.060

Haber, S. N., & Knutson, B. (2010). The reward circuit: Linking primate anatomy and human imaging. Neuropsychopharmacology, 35(1), 4–26. doi: 10.1038/npp.2009.129

Hare, T. A., O’Doherty, J., Camerer, C. F., Schultz, W., & Rangel, A. (2008). Dissociating the role of the orbitofrontal cortex and the striatum in the computation of goal values and prediction errors. The Journal of Neuroscience, 28(22), 5623–5630. doi: 10.1523/JNEUROSCI.1309-08.2008

Hayden, B. Y., Heilbronner, S. R., Pearson, J. M., & Platt, M. L. (2011). Surprise signals in anterior cingulate cortex: Neuronal encoding of unsigned reward prediction errors driving adjustment in behavior. Journal of Neuroscience, 31(11), 4178–4187. doi: 10.1523/JNEUROSCI.4652-10.2011

Jeffreys, H. (1961). *Theory of probability* (3rd ed.). Oxford, UK: Oxford University Press.

Jenkinson, M., Bannister, P., Brady, M., & Smith, S. (2002). Improved optimization for the robust and accurate linear registration and motion correction of brain images. NeuroImage, 17(2), 825–841. doi: 10.1006/nimg.2002.1132

Keuken, M. C., Bazin, P.-L., Crown, L., Hootsmans, J., Laufer, A., Müller-Axt, C., … Forstmann, B. U. (2014). Quantifying inter-individual anatomical variability in the subcortex using 7 T structural MRI. NeuroImage, 94(1), 40–46. doi: 10.1016/j.neuroimage.2014.03.032

Langkammer, C., Schweser, F., Krebs, N., Deistung, A., Goessler, W., Scheurer, E., … Reichenbach, J. R. (2012). Quantitative susceptibility mapping (QSM) as a means to measure brain iron? A post mortem validation study. NeuroImage, 62(3), 1593–1599. doi: 10.1016/j.neuroimage.2012.05.049

Logothetis, N. K., & Wandell, B. A. (2004). Interpreting the BOLD signal. Annual Review of Physiology, 66, 735–69. doi: 10.1146/annurev.physiol.66.082602.092845

Markowitz, H. (1952). Portfolio selection. The Journal of Finance, 7(1), 77–91. doi: 10.1111/j.1540-6261.1952.tb01525.x

Matsumoto, M., & Hikosaka, O. (2009). Two types of dopamine neuron distinctly convey positive and negative motivational signals. Nature, 459(7248), 837–841. doi: 10.1038/nature08028

Metere, R., Kober, T., Möller, H. E., & Schäfer, A. (2017). Simultaneous quantitative MRI mapping of T1, T∗2 and magnetic susceptibility with multi-echo MP2RAGE. PloS One, 12(1), 1–28. doi: 10.1371/journal.pone.0169265

O’Doherty, J. P., & Bossaerts, P. (2008). Toward a mechanistic understanding of human decision making: Contributions of functional neuroimaging. Current Directions in Psychological Science, 17(2), 119–123. doi: 10.1111/j.1467-8721.2008.00560.x

Pauli, W. M., Larsen, T., Collette, S., Tyszka, J. M., Seymour, B., & O’Doherty, J. P. (2015). Distinct contributions of ventromedial and dorsolateral subregions of the human substantia nigra to appetitive and aversive learning. The Journal of Neuroscience, 13(42), 14220–14233. doi: 10.1523/JNEUROSCI.2277-15.2015

Pauli, W. M., Nili, A. N., & Tyszka, J. M. (2018). A high-resolution probabilistic in vivo atlas of human subcortical brain nuclei. Scientific Data, 5(180063), 1–13. doi: 10.1038/sdata.2018.63

Peirce, J. W. (2007). PsychoPy—psychophysics software in Python. Journal of Neuroscience Methods, 162(1-2), 8–13. doi: 10.1016/j.jneumeth.2006.11.017

Power, J. D., Mitra, A., Laumann, T. O., Z.Snyder, A., L.Schlaggar, B., & E.Petersen, S. (2014). Methods to detect, characterize, and remove motion artifact in resting state fMRI. NeuroImage, 84(1), 320–341. doi: 10.1016/j.neuroimage.2013.08.048

Preuschoff, K., Bossaerts, P., & Quartz, S. R. (2006). Neural differentiation of expected reward and risk in human subcortical structures. Neuron, 51, 381–390. doi: 10.1016/j.neuron.2006.06.024

Roeper, J. (2013). Dissecting the diversity of midbrain dopamine neurons. Trends in neurosciences, 36(6), 336–342.

Schoenbaum, G., Takahashi, Y., Liu, T.-L., & McDannald, M. A. (2011). Does the orbitofrontal cortex signal value? Annals of the New York Academy of Sciences, 1239, 87–99. doi: 10.1111/j.1749-6632.2011.06210.x

Schultz, W. (1998). Predictive reward signal of dopamine neurons. Journal of neurophysiology, 80(1), 1–27. doi: 10.1152/jn.1998.80.1.1

Schultz, W. (2015). Neuronal reward and decision signals: From theories to data. Physiological Reviews, 95, 853–951. doi: 10.1152/physrev.00023.2014

Seabold, S., & Perktold, J. (2010). Statsmodels: Econometric and statistical modeling with Python. In Proceedings of the 9th Python in science conference (pp. 57–61).

Singer, T., Critchley, H. D., & Preuschoff, K. (2009). A common role of insula in feelings, empathy and uncertainty. Trends in Cognitive Sciences, 13(8), 334–340. doi: 10.1016/j.tics.2009.05.001

Sutton, R. S., & Barto, A. G. (1998). Reinforcement learning: An introduction. Cambridge: MIT press.

Trutti, A. C., Mulder, M. J., Hommel, B., & Forstmann, B. U. (2019). Functional neuroanatomical review of the ventral tegmental area. NeuroImage.

Tustison, N. J., Avants, B. B., Cook, P. A., Zheng, Y., Egan, A., Yushkevich, P. A., & Gee, J. C. (2010). N4ITK: improved N3 bias correction. IEEE Transactions on Medical Imaging, 29(6), 1310–1320. doi: 10.1109/TMI.2010.2046908

van der Zwaag, W., Schäfer, A., Marques, J. P., Turner, R., & Trampel, R. (2015). Recent applications of UHF-MRI in the study of human brain function and structure: A review. NMR in Biomedicine, 29(9), 1274–1288. doi: 10.1002/nbm.3275

Watabe-Uchida, M., Eshel, N., & Uchida, N. (2017). Neural circuitry of reward prediction error. Annal Review of Neuroscience, 40, 373–394. doi: 10.1146/annurev-neuro-072116-031109

Zaghloul, K. A., Blanco, J. A., Weidemann, C. T., McGill, K., Jaggi, J. L., Baltuch, G. H., & Kahana, M. J. (2009). Human substantia nigra neurons encode unexpected financial rewards. Science, 323(5920), 1496–1499. doi: 10.1126/science.1167342

Zhang, Y., Larcher, K., Misic, B., & Dagher, A. (2001). Segmentation of brain MR images through a hidden Markov random field model and the expectation-maximization algorithm. IEEE Transactions on Medical Imaging, 20(1), 45–57. doi: 10.1109/42.906424

Zhang, Y., Larcher, K., Misic, B., & Dagher, A. (2017). Anatomical and functional organization of the human substantia nigra and its connections. eLife, 6(e26653), 1–23. doi: 10.7554/eLife.26653

