## Supplementary Materials for "The role of dopaminergic nuclei in predicting and experiencing gains and losses: A 7T human fMRI study"

Birte U. Forstmann*
University of Amsterdam

*shared senior authorship

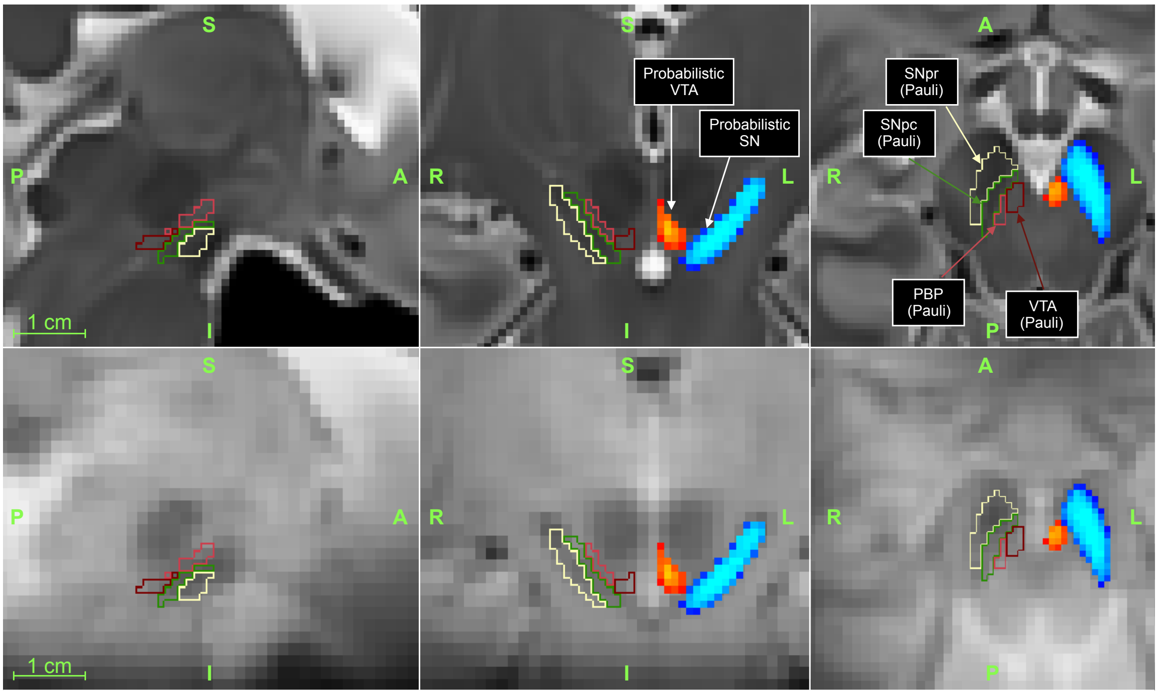


*Figure S1*. Hard lines: Pauli, Nili, and Tyszka (2018)'s deterministic masks for the ventral tegmental area (VTA), parabrachial pigmented nucleus (PBP) – which together form what is usually referred to as VTA – substantia nigra pars reticulata (SNpr), and and pars compacta(SNpc) – which together form the SN. Red to yellow gradients: probabilistic map of the VTA, estimated in the current study. Blue to light-blue gradient: probabilistic map of the SN, estimated in the current study. Note that, in the probabilistic maps, darker colors represent lower probability for a voxel to belong to the ROI. Only voxels with probability higher than 30% were kept. In the top row, the background image is Pauli et al. (2018)'s template, while the mean functional image of the current study, in MNI space, is the background in the bottom row.

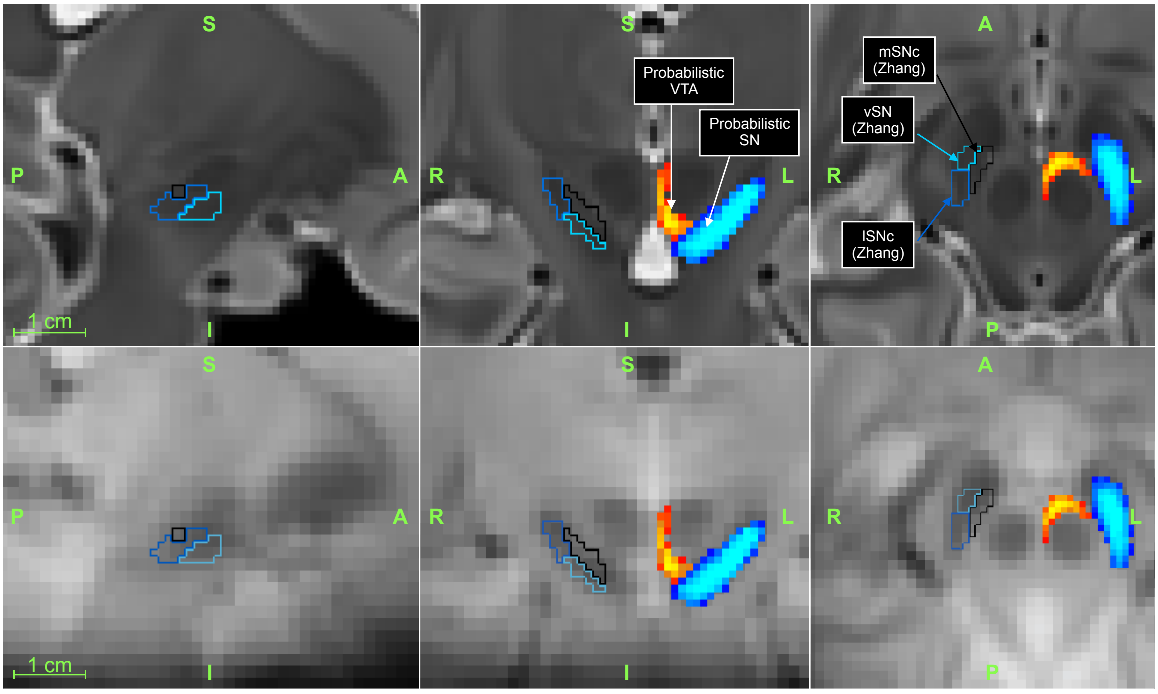


*Figure S2.* Hard lines: Zhang, Larcher, Misic, and Dagher (2017)'s deterministic masks for the medial and lateral parts of the substantia nigra pars compacta (mSNc and lSNc) and for the ventral part of the substantia nigra (vSN)– which together form the SN. Red to yellow gradients: probabilistic map of the VTA, estimated in the current study. Blue to light-blue gradient: probabilistic map of the SN, estimated in the current study. Note that, in the probabilistic maps, darker colors represent lower probability for a voxel to belong to the ROI. Only voxels with probability higher than 30% were kept. In the top row, the background image is Pauli et al. (2018)'s template, while the mean functional image of the current study, in MNI space, is the background in the bottom row.

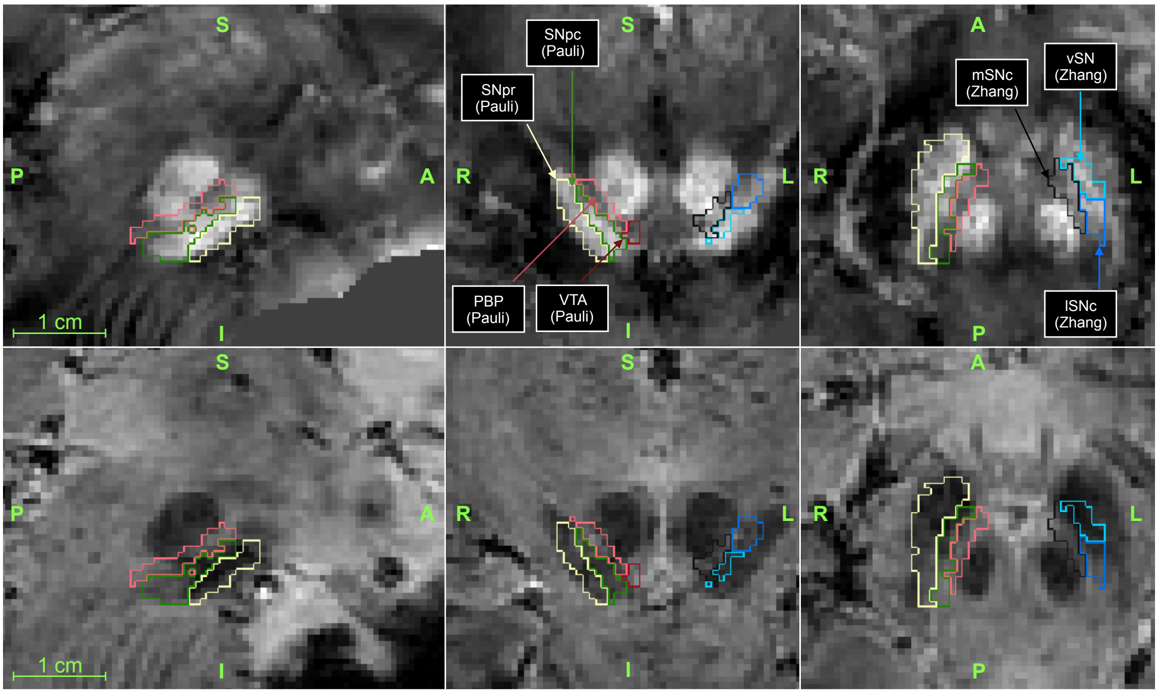


*Figure S3*. Zhang et al. (2017)'s and Pauli et al. (2018)'s deterministic masks in one participant's individual space. The image on the top row is the quantitative susceptibility mapping (QSM) and the image on the bottom is the mean of the third and fourth echo of the $T_2^*$-weighted image.

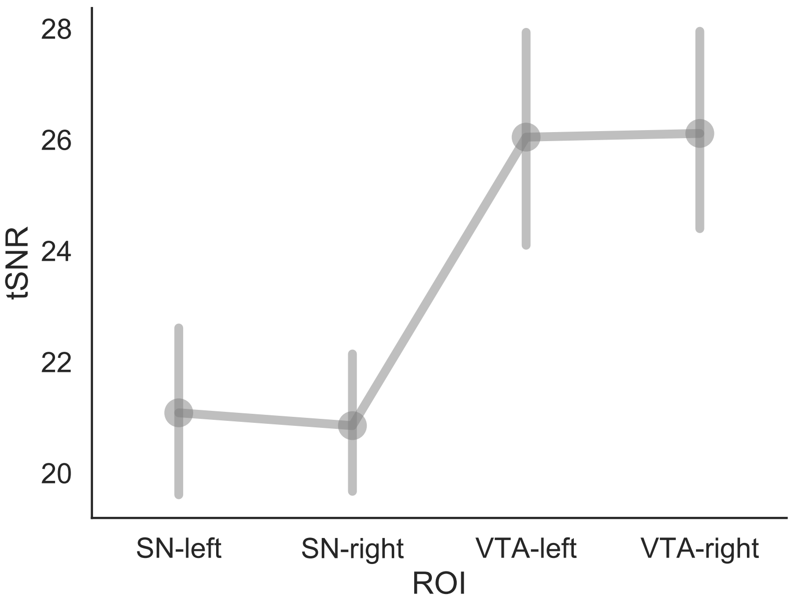


*Figure S4.* Temporal signal-to-noise ratio (tSNR) across regions of interests (ROI): left and right substantia nigra (SN) and left and right ventral tegmental area (VTA). Bars represent 95% confidence intervals.

Table S1

*Automatic labeling based on peak coordinates of the voxelwise-GLM.*

| | Predictor | Cluster Index | Voxels | Harvard-Oxford Cortical Structural Atlas | Harvard-Oxford Subcortical Structural Atlas |
|---|---|---|---|---|---|
| 0 | EV (positive) | 5 | 871 | 37% Precuneous Cortex | 40% Right Cerebral Cortex |
| 1 | EV (positive) | 4 | 868 | 63% Frontal Medial Cortex | 72% Left Cerebral Cortex |
| 2 | EV (positive) | 3 | 283 | 21% Subcallosal Cortex | 24% Left Accumbens |
| 3 | EV (positive) | 2 | 275 | * | 88% Right Accumbens, 4% Right Caudate |
| 4 | EV (positive) | 1 | 237 | 40% Frontal Pole | 46% Left Cerebral Cortex |
| 5 | EV (negative) | 3 | 1274 | * | 35% Brain-Stem, 3% Right Thalamus |
| 6 | EV (negative) | 2 | 226 | 42% Lingual Gyrus, 29% Occipital Fusiform Gyrus | 72% Left Cerebral Cortex |
| 7 | EV (negative) | 1 | 218 | 14% Intracalcarine Cortex | 26% Left Cerebral Cortex |
| 8 | risk (positive) | 5 | 7344 | * | * |
| 9 | risk (positive) | 4 | 1054 | 45% Planum Temporale, 30% Parietal Operculum C... | 87% Right Cerebral Cortex |
| 10 | risk (positive) | 3 | 591 | * | * |
| 11 | risk (positive) | 2 | 365 | * | * |
| 12 | risk (positive) | 1 | 305 | 33% Middle Temporal Gyrus, temporooccipital part | 77% Right Cerebral Cortex |
| 13 | risk (negative) | 7 | 2664 | 64% Frontal Orbital Cortex | 79% Right Cerebral Cortex |
| 14 | risk (negative) | 6 | 1300 | 63% Frontal Pole | 62% Right Cerebral Cortex |
| 15 | risk (negative) | 5 | 1209 | 56% Occipital Pole | 74% Right Cerebral Cortex |
| 16 | risk (negative) | 4 | 950 | 74% Frontal Orbital Cortex | 96% Left Cerebral Cortex |
| 17 | risk (negative) | 3 | 515 | 73% Frontal Pole | 73% Left Cerebral Cortex |
| 18 | risk (negative) | 2 | 467 | 69% Middle Temporal Gyrus, posterior division | 90% Right Cerebral Cortex |
| 19 | risk (negative) | 1 | 397 | 20% Cingulate Gyrus, anterior division | 27% Left Cerebral Cortex |
| 20 | RPE (positive) | 8 | 4234 | * | 81% Right Accumbens, 11% Right Caudate |
| 21 | RPE (positive) | 7 | 2724 | 9% Subcallosal Cortex | 3% Left Accumbens |
| 22 | RPE (positive) | 6 | 526 | * | * |
| 23 | RPE (positive) | 5 | 506 | 79% Frontal Pole | 81% Right Cerebral Cortex |
| 24 | RPE (positive) | 4 | 398 | 64% Frontal Pole | 66% Left Cerebral Cortex |
| 25 | RPE (positive) | 3 | 390 | * | 47% Brain-Stem |
| 26 | RPE (positive) | 2 | 314 | 52% Supramarginal Gyrus, anterior division | 86% Right Cerebral Cortex |
| 27 | RPE (positive) | 1 | 273 | 41% Precuneous Cortex | 45% Right Cerebral Cortex |
| 28 | surprise (positive) | 4 | 1026 | 36% Inferior Frontal Gyrus, pars triangularis | 73% Right Cerebral Cortex |
| 29 | surprise (positive) | 3 | 871 | 39% Superior Temporal Gyrus, posterior division | 91% Right Cerebral Cortex |
| 30 | surprise (positive) | 2 | 464 | 48% Middle Temporal Gyrus, posterior division | 89% Left Cerebral Cortex |
| 31 | surprise (positive) | 1 | 328 | 79% Temporal Pole | 88% Right Cerebral Cortex |
| 32 | surprise (negative) | 3 | 1291 | 33% Precuneous Cortex | 45% Left Cerebral Cortex |
| 33 | surprise (negative) | 2 | 635 | 41% Precuneous Cortex | 51% Right Cerebral Cortex |
| 34 | surprise (negative) | 1 | 250 | 30% Central Opercular Cortex | 34% Right Cerebral Cortex |

*Note.* Automatic labeling from the Harvard-Oxford Cortical and Subcortical Structural Atlases for the clusters resulting from the voxelwise general linear model (GLM). The labels are uniquely based on the peak coordinates of each cluster.

Table S2

*ROI-wise GLM results using Preuschoff, Bossaerts, and Quartz (2006)'s design.*

| ROI | EV(1st epoch) | risk (1st epoch) | EV(2nd epoch) | risk (2nd epoch) | RPE | surprise |
|---|---|---|---|---|---|---|
| SN-left | $t(26)=0.06$, $p=0.95$ | $t(26)=3.93$, $p=0.001$* | $t(26)=-0.40$, $p=0.69$ | $t(26)=-3.12$, $p=0.004$* | $t(26)=1.88$, $p=0.07$ | $t(26)=0.20$, $p=0.84$ |
| SN-right | $t(26)=1.08$, $p=0.29$ | $t(26)=2.43$, $p=0.022$* | $t(26)=-2.11$, $p=0.04$* | $t(26)=-2.86$, $p=0.008$* | $t(26)=2.33$, $p=0.03$* | $t(26)=1.55$, $p=0.13$ |
| VTA-left | $t(26)=0.75$, $p=0.46$ | $t(26)=3.56$, $p=0.001$* | $t(26)=-2.20$, $p=0.04$* | $t(26)=-2.00$, $p=0.056$ | $t(26)=2.93$, $p=0.01$* | $t(26)=-0.17$, $p=0.87$ |
| VTA-right | $t(26)=0.79$, $p=0.44$ | $t(26)=2.41$, $p=0.023$* | $t(26)=-0.92$, $p=0.37$ | $t(26)=-3.33$, $p=0.003$* | $t(26)=2.47$, $p=0.02$* | $t(26)=0.49$, $p=0.63$ |

*Note.* Results of the independent two-sided t-tests for the mean of the predictors of main interest of the GLM being equal to zero: expected value (EV) and expected risk (estimated when the trials' first number is presented), and reward prediction error (RPE) and surprise (estimated when the trial's reward or punishment are presented). As in Preuschoff et al. (2006), EV and risk were estimated separately for the 1st and 2nd epoch of the period following the presentation of the first number. These tests were run separately by regions of interest: left and right substantia nigra (SN), and left and right ventral tegmental area (VTA).

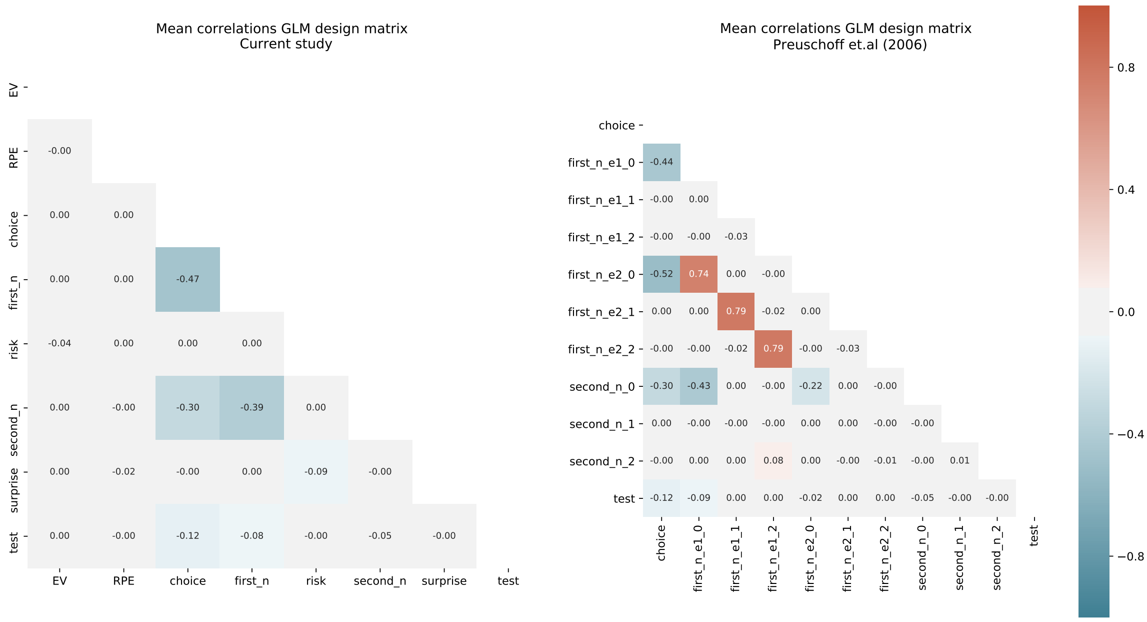


*Figure S5*. Correlation across the main predictors in the general linear model (GLM) as set up in the main analyses in this study (on the right) and in the control analyses (on the left), in which the GLM is set up similarly to Preuschoff et al. (2006). In these analyses, the intercepts (first__n__e1__0 and first__n__e2__0), first order (first__n__e1__1 and first__n__e2__1) and second order (first__n__e1__2 and first__n__e2__2) regressors for the first number are highly correlated across the two epochs in which this event is divided (e1 and e2). Correlations were first computed at the run level, separately for each subject, and then averaged across runs and subjects.

### References

Pauli, W. M., Nili, A. N., & Tyszka, J. M. (2018). A high-resolution probabilistic in vivo atlas of human subcortical brain nuclei. *Scientific Data*, *5*(180063), 1–13. doi: 10.1038/sdata.2018.63

Preuschoff, K., Bossaerts, P., & Quartz, S. R. (2006). Neural differentiation of expected reward and risk in human subcortical structures. *Neuron*, *51*, 381–390. doi: 10.1016/j.neuron.2006.06.024

Zhang, Y., Larcher, K., Misic, B., & Dagher, A. (2017). Anatomical and functional organization of the human substantia nigra and its connections. *eLife*, *6*(e26653), 1–23. doi: 10.7554/eLife.26653
